# Targeting CTP Synthetase 1 to Restore Interferon Induction and Impede Nucleotide Synthesis in SARS-CoV-2 Infection

**DOI:** 10.1101/2021.02.05.429959

**Authors:** Youliang Rao, Ting-Yu Wang, Chao Qin, Bianca Espinosa, Qizhi Liu, Arunika Ekanayake, Jun Zhao, Ali Can Savas, Shu Zhang, Mehrnaz Zarinfar, Yongzhen Liu, Wenjie Zhu, Nicholas Graham, Taijiao Jiang, Chao Zhang, Pinghui Feng

## Abstract

The newly emerged SARS-CoV-2 caused a global pandemic with astonishing mortality and morbidity. The mechanisms underpinning its highly infectious nature remain poorly understood. We report here that SARS-CoV-2 exploits cellular CTP synthetase 1 (CTPS1) to promote CTP synthesis and suppress interferon (IFN) induction. Screening a SARS-CoV-2 expression library identified ORF7b and ORF8 that suppressed IFN induction via inducing the deamidation of interferon regulatory factor 3 (IRF3). Deamidated IRF3 fails to bind the promoters of classic IRF3-responsible genes, thus muting IFN induction. Conversely, a shRNA-mediated screen focused on cellular glutamine amidotransferases corroborated that CTPS1 deamidates IRF3 to inhibit IFN induction. Functionally, ORF7b and ORF8 activate CTPS1 to promote *de novo* CTP synthesis while shutting down IFN induction. *De novo* synthesis of small-molecule inhibitors of CTPS1 enabled CTP depletion and IFN induction in SARS-CoV-2 infection, thus impeding SARS-CoV-2 replication. Our work uncovers a strategy that a viral pathogen couples immune evasion to metabolic activation to fuel viral replication. Inhibition of the cellular CTPS1 offers an attractive means for developing antiviral therapy that would be resistant to SARS-CoV-2 mutation.

## INTRODUCTION

First reported in December 2019, the coronavirus disease known as COVID-19 rapidly spread worldwide and became a global pandemic, causing more than 100 million infections and claiming more than 2.16 million lives by January 2021. The etiological agent of COVID-19 was soon identified as a new coronavirus, severe acute respiratory syndrome coronavirus 2 (SARS-CoV-2)(Hu et al., 2020; Zhou et al., 2020). Despite the rapid development of effective vaccine against SARS-CoV-2, COVID-19 patients urgently require post-infection treatment or therapeutics (Dong et al., 2020; Krammer, 2020). Current management of severe COVID-19 patients primarily consists of supportive care that optimizes oxygen administration via intubation and mechanical ventilation (Dondorp et al., 2020). Treatment options are limited to repurposed drugs, such as remdesivir (Beigel et al., 2020; Goldman et al., 2020; Pruijssers et al., 2020; Wang et al., 2020b) and dexamethasone (Group et al., 2020), and their clinical effect on severe or critical COVID-19 patients is yet to be established with well controlled trials.

Compared to previous zoonotic coronaviruses including SARS-CoV and MERS-CoV, SARS-CoV-2 is highly infectious and transmissible (Harrison et al., 2020; Mona Fani et al., 2020). Current efforts have extensively focused on the entry step, which is primarily mediated by the interaction between SARS-CoV-2 spike (S) protein and the human angiotensin-converting enzyme 2 (hACE2) (Hoffmann et al., 2020; Zhou et al., 2020). Structural and functional analyses of this receptor-ligand interaction indicate that SARS-CoV-2 S protein evolved higher affinity for binding to hACE2 on target cells, partly explaining the highly infectious nature of SARS-CoV-2 during COVID-19 pandemic (Wang et al., 2020a). However, little is known about the viral mechanisms downstream of viral entry that contribute to the infection and pathogenesis of SARS-CoV-2. Innate immune response constitutes the first line of defense against intracellular pathogens such as viruses (Takeuchi and Akira, 2010). To efficiently replicate within an immune-competent host, a virus must overcome the barrier of host innate immune defense, chiefly mediated by the interferon system (Nelemans and Kikkert, 2019). Indeed, previous studies of SARS-CoV-2 infection involving patient samples, model animals and cell lines suggest that SARS-CoV-2 either weakly induces IFNs or inhibits IFN induction (Lei et al., 2020; O’Brien et al., 2020; Park and Iwasaki, 2020). How exactly SARS-CoV-2 interacts with the cellular IFN system remains largely unknown, except for sequence analysis comparing SARS-CoV-2 to SARS-CoV and MERS-CoV that was used to predict plausible functions of viral proteins (Gordon et al., 2020).

In addition to overcoming innate immune defense, viruses rely on cellular machinery to synthesize macromolecules and biomaterials that are subsequently assembled into progeny virions (de Castro et al., 2013; Thaker et al., 2019). Thus, viruses often activate and redirect cellular biosynthetic activities to facilitate the production of viral components, such as proteins, nucleic acids and lipids, that constitute essential building blocks of virions (Eisenreich et al., 2019; Gita Mahmoudabadi et al., 2017). Central to viral replication is the reprogramming of cellular metabolic processes that are often activated to provide precursors for viral biosynthesis in infected cells (Thaker et al., 2019). The highly infectious nature of SARS-CoV-2 likely involves molecular interactions that boost the rate-limiting steps of key metabolic pathways to fuel viral replication and subsequent dissemination (Ayres, 2020; Pislar et al., 2020). Cellular glutamine amidotransferases (GATs) catalyze the synthesis of nucleotides, amino acids, glycoproteins and an enzyme cofactor (NAD) (F. Massière and Badet-Denisot., 1998; Walker and van der Donk, 2016). Our studies have shown that these enzymes are capable of deamidating key signaling molecules, such as those involved in innate immune defense, to modulate fundamental biological processes (Zhao et al., 2016a). Recent studies from our group suggest that cellular GATs potentially couple innate immune response to cellular metabolism via deamidating checkpoint proteins, e.g., RIG-I and RelA (He et al., 2015; Zhao et al., 2020; Zhao et al., 2016b).

To probe the inhibition of IFN response by SARS-CoV-2, we identified viral-host interactions that couple inhibition of IFN induction to activated CTP synthesis during SARS-CoV-2 infection. Specifically, SARS-CoV-2 ORF8 and, to a lesser extent, ORF7b interact with and activate CTP synthetase 1 (CTPS1) to promote CTP synthesis. Activated CTPS1 also deamidates IRF3 to inhibit IFN induction and downstream immune response. As such, several small-molecule that were developed as CTPS1 inhibitors restored IFN induction and depleted CTP in SARS-CoV-2-infected cells, thus impeding SARS-CoV-2 replication. This study unravels a viral strategy that hijacks a cellular CTP synthesis enzyme to fuel nucleotide synthesis and shut down IFN induction, forging a molecular link between metabolism and innate immune defense. CTPS1 blockade depletes CTP supply and restores innate immune response, which can yield antiviral therapy that is resistant to SARS-CoV-2 genetic variation.

## RESULTS

### Identification of SARS-CoV-2 Proteins that Induce IRF3 Deamidation and Inhibit IFN Induction

SARS-CoV-2 is highly infectious and transmissible in human population. Ongoing research is keen on viral entry, how viral post-entry mechanisms contribute to the SARS-CoV-2 infection is poorly understood. We hypothesized that SARS-CoV-2 encodes a number of viral polypeptides to modulate host innate immune response, thereby promoting viral replication and dissemination. To test this hypothesis, we first compared the antiviral gene expression induced by SARS-CoV-2 with that induced by Sendai virus, a prototype RNA virus. In normal human bronchial epithelial cells (NHBE), Sendai virus triggered a rapid and robust expression of multiple antiviral genes with a peak at 6 hour post-infection (hpi) and a fold increase ranging from ~15,000 (for *ISG15*) to 800,000 (for *IFNB1*) (Figure 1A). By stark contrast, SARS-CoV-2 induced a weak and delayed expression of antiviral genes, peaking at 96 hpi (Figure 1A). The fold induction of these antiviral genes by SARS-CoV-2 was roughly three orders of magnitude lower than that induced by Sendai virus. Similar patterns were observed in human Calu-3 lung cancer cells and Caco-2 colorectal cancer cells, two cell lines that support robust SARS-CoV-2 replication (Figure S1A and S1B). Interestingly, although delayed, the expression of *Mx1* in Calu-3 and Caco-2 cells induced by SARS-CoV-2 was as robust as that induced by Sendai virus. To determine whether RNA derived from SARS-CoV-2-infected cells is able to provoke innate immune activation, we extracted total RNA from SARS-CoV-2-infected NHBE cells, and along with poly(I:C), transfected it into NHBE cells. When mRNA of *IFNB1*, *ISG15*, *ISG56*, *CCL5* and *Mx1* was analyzed, we found that the total RNA extracted from SARS-CoV-2-infected NHBE cells, but not from mock-infected NHBE cells, induced antiviral gene expression as potently as poly(I:C) (Figure 1B). These results suggest that SARS-CoV-2 is able to suppress antiviral innate immune defense.

**Figure 1.**
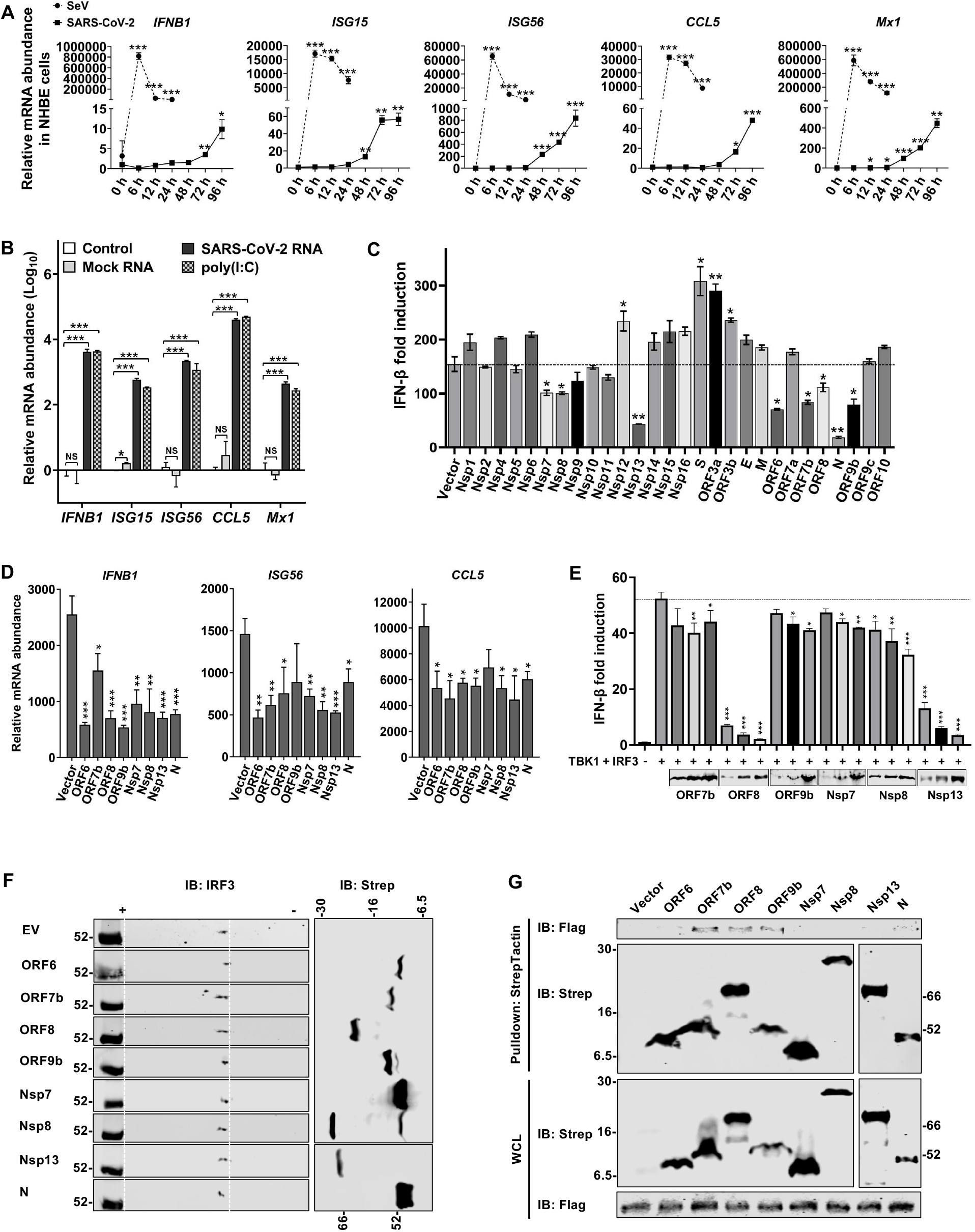
Identification of SARS-CoV-2 proteins that induce IRF3 deamidation to inhibit IFN induction. (A) NHBE cells were infected with Sendai virus (SeV) (100 HAU/ml) or SARS-CoV-2 (MOI = 1). Total RNA was extracted, reverse transcribed and analyzed by real-time PCR with primers specific for *IFNB1*, *ISG15*, *ISG56*, *CCL5* and *Mx1*. (B) NHBE cells were transfected with poly(I:C) and RNA isolated from mock- or SARS-CoV-2-infected NHBE cells (MOI = 1, 72 hpi). RNA extraction and real-time PCR were performed as in (A). (C) Modulation of IFN-β induction was determined by a promoter activity in 293T cells expressing indicated SARS-CoV-2 proteins, with SeV infection. (D) Inhibition of antiviral gene expression by SARS-CoV-2 proteins in 293T cells infected with SeV was examined by real-time PCR with primers specific for indicated genes. (E) Inhibition of IFN-β induction by selected SARS-CoV-2 proteins was determined by reporter assay of 293T cells expressing TBK-1 and IRF3. (F) Effect of selected SARS-CoV-2 proteins on IRF3 charge status was determined by two-dimensional gel electrophoresis and immunoblotting analyses using lysates of 293T cells transfected with plasmids containing indicated genes. (G) Interactions between exogenous IRF3 and SARS-CoV-2 proteins were analyzed by co-immunoprecipitation in transfected 293T cells. For (F) and (G), Strep indicates SARS-CoV-2 proteins. Error bars indicate standard deviation (SD) of technical triplicates. Statistical significance was calculated using unpaired, two-tailed Student’s *t*-test. \**P* < 0.05; \*\**P* < 0.01; \*\*\**P* < 0.001. See related **Figure S1**.

To understand the viral mechanisms of immune modulation, we screened viral polypeptides with an expression library for IFN-β induction by Sendai virus. This reporter assay identified several SARS-CoV-2 proteins, including ORF6, ORF7b, ORF8, ORF9b, N, Nsp7, Nsp8 and Nsp13, that inhibit IFN induction to various extent (Figure 1C). To validate, we transfected 293T cells with plasmids expressing these individual viral polypeptides and found that all eight viral polypeptides inhibited the expression of antiviral genes, including *IFNB1*, *ISG56* and *CCL5*, in response to Sendai virus infection (Figure 1D). Importantly, ORF6 and N were previously reported to inhibit the nuclear import of transcription factors (e.g., IRF3) and sequester viral double-stranded RNA, respectively (Cascarina and Ross, 2020; Xia et al., 2020), to suppress innate immune defense. Thus, we further examined the other six viral proteins for the inhibition of IFN induction. Nevertheless, these results demonstrate that multiple viral polypeptides can inhibit IFN induction.

SARS-CoV-2 is a coronavirus that likely induces innate immune activation via RNA sensors such as RIG-I and MDA5. To determine the target of inhibition, we over-expressed key components of this pathway, including RIG-I-N (2CARD-only), MAVS, TBK1 and IRF3, and examined IFN-β induction by reporter assay (Figure S1C). Notably, RIG-I-N is a constitutively active form of RIG-I independent of RNA ligand. This assay showed that ORF7b, ORF8 and Nsp13 can significantly inhibit IFN induction by the expression of more than one component of the RIG-I-IFN pathway, while ORF9b, Nsp7 and Nsp8 did not significantly inhibit IFN induction by ectopic expression (Figure S1D). Further analysis demonstrated that ORF8 and Nsp13 potently inhibited IFN induction by over-expressing IRF3 and TBK1, suggesting that IRF3 is the point of inhibition (Figure 1E). Thus, we focused on IRF3 regulation by these viral polypeptides, with a keen interest in deamidation, a process that can be catalyzed by metabolic glutamine amidotransferases (Zhao et al., 2016a). When endogenous IRF3 was analyzed by two-dimensional gel electrophoresis, we found that ORF7b, ORF8, NSP8 and NSP13 induced a shift of IRF3 toward the negative side of the gel strip, suggesting the deamidation of IRF3 (Figure 1F). Interestingly, expression of N induced a shift of IRF3 in the upper-left direction, suggesting possible phosphorylation. We also investigated interaction between IRF3 and these viral proteins by co-immunoprecipitation (Co-IP). As shown in Figure 1G, IRF3 interacted with ORF7b, ORF8, and ORF9b, but not ORF6, Nsp7, Nsp8 or Nsp13.These results collectively show that SARS-CoV-2 targets IRF3 for inhibition.

### CTPS1 Inhibits IFN Induction

Given that multiple SARS-CoV-2 viral polypeptides induce IRF3 deamidation, we reasoned that a cellular enzyme catalyzes IRF3 deamidation. The human genome encodes 11 glutamine amidotransferases that can potentially function as protein deamidases (Li et al., 2019; Zhao et al., 2020). With shRNA-mediated knockdown, we screened for cellular GATs, when knocked down, increase IFN induction by Sendai virus infection. This experiment showed that depletion of CTPS1 elevated Sendai virus-induced IFN expression by ~1.5-fold (Figure 2A). Remarkably, depletion of the closely-related CTPS2 had no significant effect on IFN induction. The knockdown efficiency of these cellular GATs was validated in our recent publication (Li et al., 2019). Notably, depletion of several GATs, including PPAT, ASNS and NADSYN1, reduced IFN induction, possibly due to the essential roles of these metabolic enzymes in cell proliferation and survival. To examine the role of CTPS1 in innate immune defense, we constructed two 293T cell lines that each expresses a unique shRNA and validated CTPS1 depletion by quantitative real-time PCR and immunoblotting (Figure 2B and S2A). Compared with control cells, Sendai virus infection induced higher transcript levels of *IFNB1*, *CCL5*, *ISG56* and *ISG15*, in CTPS1-depleted 293T cells, as analyzed by real-time PCR (Figure 2C). The elevated production of IFN-β and CCL5 was further confirmed by ELISA (Figure 2D). Similarly, depletion of CTPS1 in human THP-1 monocytes (Figure 2E) elevated IFN-β and CCL5 expression and production in response to Sendai virus infection (Figure 2F and S2B). Consistent with the elevated antiviral immune response, depletion of CTPS1 reduced the replication of vesicular stomatitis virus (VSV) by ~10-fold in 293T cells (Figure 2G). These results collectively demonstrate that CTPS1 negatively regulates IFN induction in response to Sendai virus infection.

**Figure 2.**
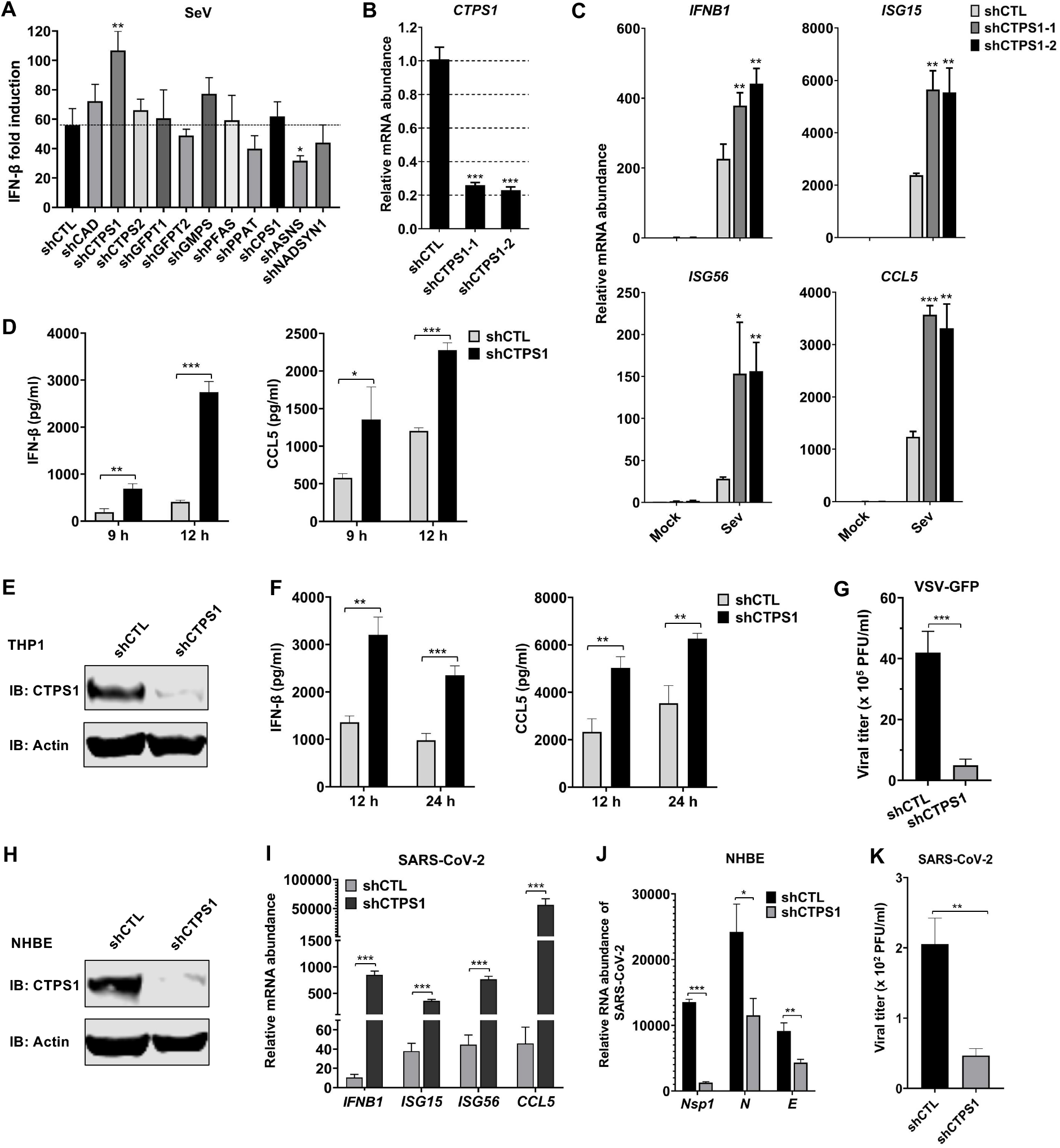
CTPS1 inhibits IFN induction. (A) Effect of cellular glutamine amidotransferases (GATs) on IFN induction by Sendai virus infection was determined by luciferase assay using GAT-depleted 293T cells transfected with the IFN-β reporter cocktail: CAD: carbamoyl-phosphate synthetase, aspartate transcarbomylase, and dihydroorotase; CTPS: CTP synthetase; GFPT: glutamine fructose-6-phosphate amidotransferase; GMPS: GMP synthetase; PFAS: phosphoribosylformylglycinamidine synthetase; PPAT: phosphoribosyl pyrophosphate amidotransferase; CPS1: carbamoyl-phosphate synthetase; ASNS: Asparagine synthetase; and NADSYN1: NAD synthetase 1. CTL, control. (B - D) Depletion of CTPS1 was validated by real-time PCR analysis using total RNA extracted from 293T cells infected with lentivirus containing indicated shRNA (B). The mRNA abundance of antiviral genes induced by Sendai virus infection was determined by real-time PCR with primers specific for indicated genes (C). Medium of Sendai virus-infected 293T cells was assessed by ELISA for IFN-β and CCL5 (D). (E and F) Knockdown of CTPS1 in THP1 monocytes was determined by immunoblotting using cells with lentivirus containing control (CTL) or CTPS1 shRNA (E). IFN-β and CCL5 in the medium of THP1 infected with Sendai virus were assessed by ELISA (F). (G) Vesicular stomatitis virus (VSV) replication in control and CTPS1-depleted 293T cells was determined by plaque assay at 10 hours post-infection (MOI=0.01). (H - K) CTPS1 depletion in NHBE cells was determined by immunoblotting (H). Effects of CTPS1 depletion on the expression of cellular antiviral genes (I) and viral genes (J) were determined by real-time PCR analysis of total RNA extracted at 48 h after SARS-CoV-2 infection (MOI =0.1). Medium of NHBE cells infected with SARS-CoV-2 was used for plaque assay to determine infectious viral progeny (K). Error bars indicate SD of technical triplicates. Statistical significance was calculated using unpaired, two-tailed Student’s *t*-test. \**P* < 0.05; \*\**P* < 0.01; \*\*\**P* < 0.001. See related **Figure S2**.

To assess the role of CTPS1 in SARS-CoV-2 infection, we depleted CTPS1 in NHBE cells (Figure 2H) and assessed antiviral gene expression and viral replication. Real-time PCR analysis indicated that depletion of CTPS1 increased the expression of *IFNB1*, *ISG15*, *ISG56* and *CCL5* by a factor ranging from ~15 (for *ISG15* and *ISG56*) to 1000 (for *CCL5*) (Figure 2I). Conversely, depletion of CTPS1 reduced SARS-CoV-2 RNA abundance by two-fold for N and E genes, and >10-fold for Nsp1 gene (Figure 2J), which correlated with a four-fold reduction in viral titer in the medium at 48 h post-infection (Figure 2K). Similar results were observed in Caco-2 cells for elevated antiviral gene expression in response to SARS-CoV-2 infection upon CTPS1 depletion (Figure S2C and S2D). This result also correlated with reduced SARS-CoV-2 replication as analyzed by real-time PCR for viral RNA abundance and plaque assay for infectious virions in the medium (Figure S2E and S2F). These results show that CTPS1 negatively regulates antiviral immune response against SARS-CoV-2 and that deficiency in CTPS1 promotes antiviral gene expression to impede SARS-CoV-2 replication.

### CTPS1 Interacts with and Deamidates IRF3

To determine the point of inhibition by CTPS1, we used an ectopic expression system as described in Figure S1C. A reporter assay indicated that depletion of CTPS1 increased the IFN-β induction activated by the ectopic expression of all components of the RIG-I-IFN pathway, including RIG-I-N, TBK1 and IRF3-5D (Figure 3A). The specific effect of CTPS1 depletion on IFN-β expression induced by IRF3-5D supports the conclusion that deamidation likely targets IRF3 for inhibition (Figure S3A). CTPS1 is a metabolic enzyme that catalyzes the synthesis of CTP from UTP and glutamine. We hypothesized that CTPS1 targets IRF3 for deamidation to inhibit IFN induction. To test this hypothesis, we first determined whether the enzyme activity of CTPS1 is required for its inhibition. We generated an enzyme-deficient mutant (C399A/H526A/E528A) of CTPS1 (CTPS1-ED) and performed a reporter assay. This result showed that wild-type CTPS1, but not the CTPS1-ED mutant, inhibited IFN-β induction induced by Sendai virus infection in a dose-dependent manner (Figure S3B). Furthermore, a non-selective inhibitor of cellular GATs, 6-diazo-5-oxo-L-norleucine (DON) elevated Sendai virus-induced IFN-β gene expression in control and CTPS1-depleted 293T cells (Figure S3C), although the effect of DON was slightly reduced when CTPS1 was depleted in 293T cells. These results support the conclusion that the enzyme activity of CTPS1 is necessary for the inhibition of IFN induction.

**Figure 3.**
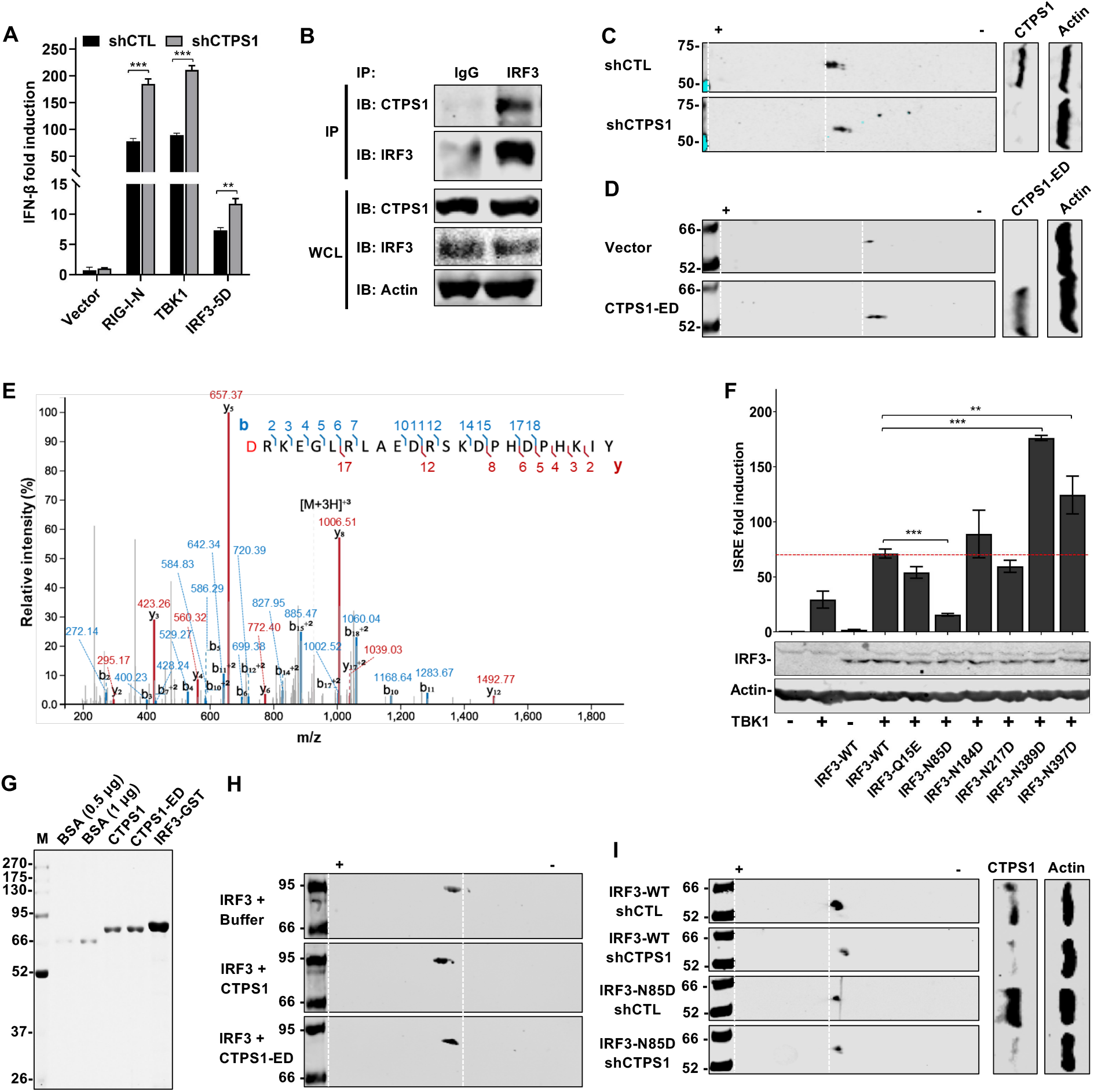
CTPS1 interacts with and deamidates IRF3. (A) Effect of CTPS1 depletion on IFN induction was determined by luciferase assay in control and CTPS1-depleted 293T cells transfected with the IFN-β reporter cocktail and plasmids containing indicated components of the RIG-I-IFN pathway. (B) Interaction between endogenous IRF3 and CTPS1 was determined by co-immunoprecipitation using 293T cell lysates and immunoblotting analyses. (C and D) Effect of CTPS1 depletion on IRF3 charge status was analyzed by two-dimensional gel electrophoresis and immunoblotting using 293T cells depleted of CTPS1 (C) or expressing the enzyme-deficient CTPS1-ED mutant (D). (E) IRF3 deamidation was determined by tandem mass spectrometry using affinity purified IRF3 in the presence of CTPS1-ED. The m/z spectrum of the peptide containing N85D is shown with the deamidated D residue highlighted in red. (F) Effects of several deamidations on IRF3 were determined by luciferase assay using 293T cells transfected with the IFN-β reporter cocktail and plasmids containing TBK-1, wild-type IRF3 or indicated deamidated IRF3 mutants. Whole cell lysates were analyzed by immunoblotting for IRF3 and TBK1 expression. (G and H) Flag-CTPS1, Flag-CTPS1-ED and IRF3-GST proteins were purified from transfected 293T cells by affinity chromatography and analyzed by Coomassie blue staining (G). BSA, bovine serum albumin. *In vitro* deamidation reactions were analyzed by two-dimensional gel electrophoresis and immunoblotting (H). (I) Effect of CTPS1 on IRF3-N85D was determined by two-dimensional gel electrophoresis and immunoblotting with control (CTL) and CTPS1-depleted 293T cells transfected with a plasmid containing Flag-IRF3-WT or Flag-IRF3-N85D. Error bars indicate SD of technical triplicates. Statistical significance was calculated using unpaired, two-tailed Student’s t-test. **P < 0.01; ***P < 0.001. See related Figure S3.

Next, we assessed whether CTPS1 interacts with IRF3 by co-immunoprecipitation (Co-IP). As shown in Figure 3B, CTPS1 was readily detected in protein complexes precipitated with antibody against IRF3, indicating the physical interaction between IRF3 and CTPS1. The interaction between IRF3 and CTPS1 was confirmed by co-IP assay from transiently transfected 293T cells, whereas CTPS2 failed to interact with IRF3 (Figure S3D). To determine whether CTPS1 is a plausible deamidase, we depleted CTPS1 with shRNA and examined IRF3 charge by two-dimensional gel electrophoresis. Indeed, knockdown of CTPS1 shifted IRF3 toward the negative end of the gel strip, indicating the increased charge due to CTPS1 depletion (Figure 3C). Similarly, the ectopic expression of the enzyme-deficient CTPS1-ED mutant also shifted IRF3 toward the negative pole of the gel strip (Figure 3D). We then sought to identify the site of deamidation using tandem mass spectrometry. We purified IRF3 in transfected 293T cells, without or with the ectopic expression of CTPS1-ED. Tandem mass spectrometry analysis consistently identified N85, N389 and N397 as deamidation sites (Figure 3E, S3E and S3F). We generated IRF3 mutants containing individual deamidated residues, i.e., Q>E and N>D mutations, and analyzed their activity in IFN induction. Remarkably, the N85D mutation nearly deprived IRF3 of the ability to activate the *IFNB1* promoter, while N389D and N397D elevated the ability of IRF3 to do so (Figure 3F and S3G). Given that the phenotype of the IRF3-N85D mutant is consistent with the inhibition of CTPS1-mediated deamidation, we focused on this mutant in the remainder of this study.

To determine whether CTPS1 is a *bona fide* deamidase of IRF3, we purified GST-IRF3, CTPS1 wild-type and the enzyme-deficient CTPS1-ED mutant from 293T cells to high homogeneity (Figure 3G) and performed IRF3 *in vitro* deamidation assays. When IRF3 was analyzed by two-dimensional gel electrophoresis, CTPS1, but not the CTPS1-ED mutant, shifted IRF3 toward the positive end of the gel strip, consistent with the increased negative charge resulting from deamidation (Figure 3H). This result indicates that CTPS1 can function as a *bona fide* deamidase of IRF3. To assess the specificity of CTPS1-mediated deamidation, we used the deamidated IRF3-N85D mutant and CTPS1 depletion for IRF3 charge analysis. Two-dimensional gel electrophoresis analysis indicated that depletion of CTPS1 shifted wild-type IRF3 toward the negative pole of the gel strip, but had no effect on the deamidated IRF3-N85D (Figure 3I). Similarly, depletion of CTPS1 failed to shift IRF3-N85A and IRF3-N85Q, indicating that IRF3-N85A and IRF3-N85Q are resistant to CTPS1-mediated deamidation (Figure S3H). Consistent with this result, IRF3-N85A demonstrated higher IFN induction in a reporter assay compared to wild-type IRF3 (Figure S3I). However, IRF3-N85Q had lower IFN induction than the wild-type, likely due to steric hindrance in DNA-binding (see next section). Together, these results support the conclusion that CTPS1 targets N85 of IRF3 for deamidation.

### Deamidated IRF3 Fails to Bind Cognate Responsive Elements within Promoters of Pro-inflammatory Genes

Having established CTPS1 negatively impacts IFN induction, we thus probed the deamidation in regulating IRF3-mediated inflammatory gene expression using the deamidated IRF3-N85D and deamidation-resistant IRF3-N85A mutants. Considering that IRF3 and IRF7 have overlapped function in inducing the expression of IFNs and antiviral genes, we “reconstituted” IRF3 expression in *Irf3*^*−/−*^*Irf7*^*−/−*^ mouse embryonic fibroblasts (MEFs) (Figure 4A) and examined antiviral immune response upon Sendai virus infection.

**Figure 4.**
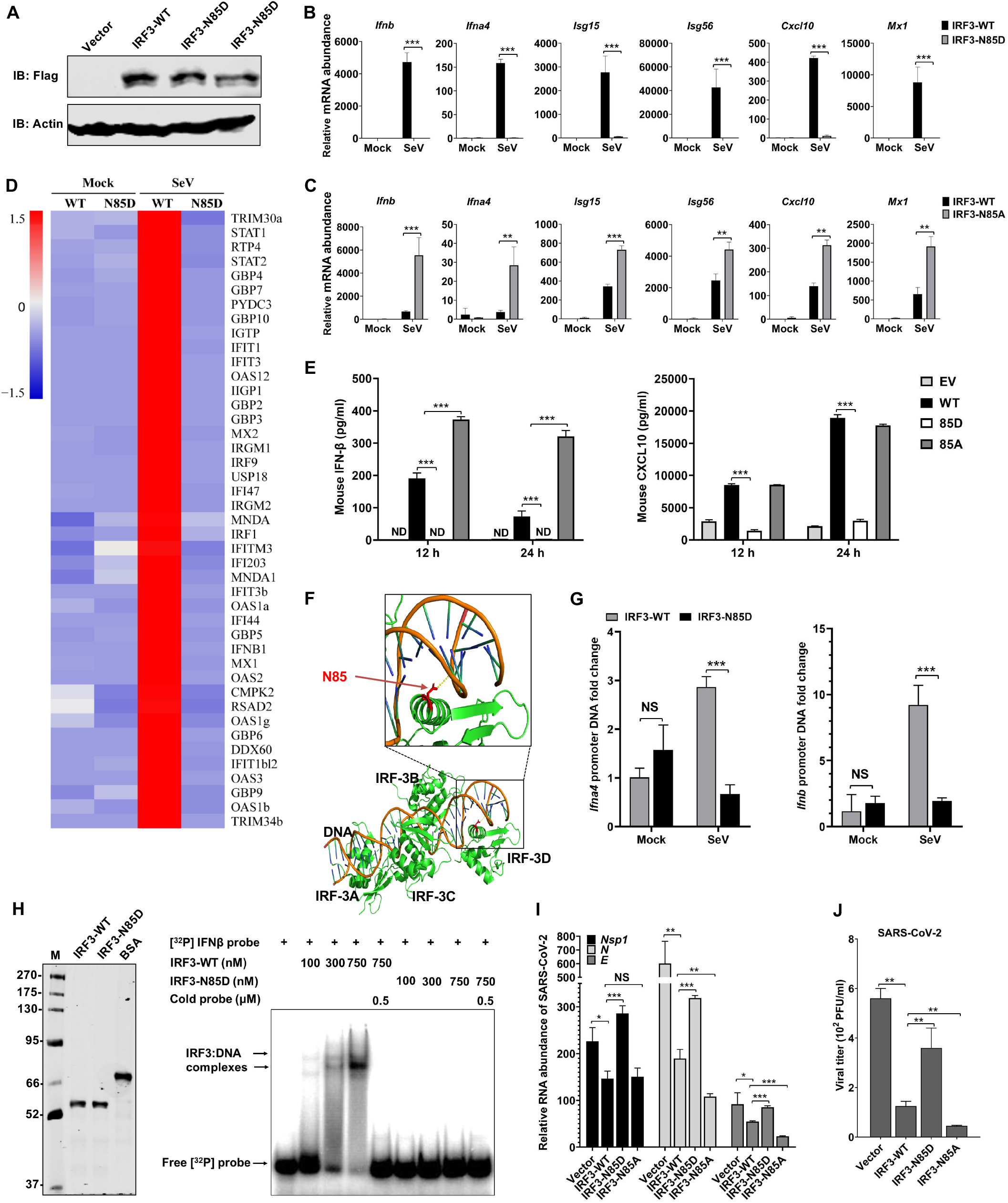
Deamidation impedes IRF3 to activate antiviral immune responses by blocking its DNA binding activity. (A) IRF3 expression was analyzed by immunoblotting in *Irf3*^*−/−*^*Irf7*^*−/−*^ MEFs reconstituted with wild-type IRF3 and its mutants. (B and C) The mRNA abundance of antiviral genes in reconstituted MEF as described in (A), with Sendai virus infection, was analyzed by real-time PCR. (D) A heatmap of the expression of IFN-related genes analyzed by RNA sequencing using total RNA extracted from Sendai virus-infected *Irf3*^*−/−*^*Irf7*^*−/−*^ MEFs reconstituted with IRF3-WT and IRF3-N85D. (E) IFN-β and CXCL10 in the medium of reconstituted *Irf3*^*−/−*^*Irf7*^*−/−*^ MEF (as described in A) infected with Sendai virus for 12 hours were assessed by ELISA. (F) Structure of the IRF3-containing *Ifnb* enhanceosome (PDB: 206G). Deamidated residue (N85) was highlighted in red. (G) Quantification by ChIP-qPCR of mouse *Ifnb* and *Ifna4* promoter sequences that were precipitated in Sendai virus-infected *Irf3*^*−/−*^*Irf7*^*−/−*^ MEFs reconstituted with IRF3-WT and IRF3-N85D. (H) IRF3-WT and IRF3-N85D proteins were purified from 293T cells by affinity chromatography and analyzed by Coomassie blue staining (left panel). BSA, bovine serum albumin. *In vitro* IRF3-DNA binding was performed and analyzed by EMSA (right panel). (I and J) Human ACE2-expressing *Irf3*^*−/−*^*Irf7*^*−/−*^ MEFs were reconstituted with IRF3-WT, IRF3-N85D, IRF3-N85A and Vector. Effect of IRF3 and its mutants on SARS-CoV-2 RNA abundance (I) was assessed by real-time PCR with total RNA extracted at 24 h after SARS-CoV-2 infection (MOI =0.01). Medium of the cells infected with SARS-CoV-2 was used for plaque assay to determine infectious viral progeny (J). Error bars indicate SD of technical triplicates. Statistical significance was calculated using unpaired, two-tailed Student’s *t*-test. \**P* < 0.05; \*\**P* < 0.01; \*\*\**P* < 0.001. See related **Figure S4**.

Real-time PCR analysis indicated that IRF3-N85D, compared to the wild-type, failed to induce the expression of IFNs and IFN-stimulated genes (Figure 4B). In contrast, the deamidation-resistant IRF3-N85A mutant more potently activated the expression of these IFNs and ISGs than wild-type IRF3 (Figure 4C). To profile the global gene expression in MEFs “reconstituted” with IRF3-N85D, we performed RNA sequencing and discovered that reconstituted expression of wild-type IRF3 activated the expression of a broad spectrum of IFNs and ISGs, while that of IRF3-N85D failed to do so (Figure 4D). The ability of IRF3 wild-type, IRF3-N85D and IRF3-N85A to activate antiviral gene expression also correlated with IFN-β and CXCL10 production in the medium in response to Sendai virus infection (Figure 4E). These results show that deamidation inhibits IRF3-mediated expression of IFNs and ISGs.

In response to Sendai virus infection, IRF3 undergoes phosphorylation, dimerization and nuclear translocation to activate the expression of inflammatory genes. To probe the effect of deamidation on IRF3 activation, we analyzed these events of IRF3 activation using wild-type IRF3 and IRF3-N85D. Immunoblotting analyses, with phosphor-specific antibody for IRF3 and native gel electrophoresis, indicated that IRF3-N85D was phosphorylated and dimerized at higher levels than wild-type IRF3 (Figure S4A). Immunofluorescence microscopy analysis showed that wild-type IRF3 and IRF3-N85D accumulated in the nucleus at similar rates (Figure S4B and S4C). Thus, deamidation did not impair the phosphorylation, dimerization or nuclear translocation of IRF3 when activated by Sendai virus infection.

In collaboration with other transcription factors, IRF3 acts in concert in the so-called “enhanceosome” of the IFN-β promoter, which has been well characterized by structural studies (Panne et al., 2007). In the structure of the DNA-protein complex (Panne et al., 2007), the N85 residue of IRF3 makes direct contact with the backbone of dsDNA via hydrogen bond (Figure 4F). Deamidation of N85 is predicted to disrupt the hydrogen bond and create a negative charge that likely repels the highly negatively charged backbone of dsDNA. Thus, we performed chromosome immunoprecipitation and quantified DNA by real-time PCR. Wild-type IRF3, but not IRF3-N85D, enriched the sequences of IFNs, including *Ifnb* and *Ifna4*, in response to Sendai virus infection (Figure 4G). Consistent with this, an *in vitro* gel shift assay using purified IRF3 proteins indicated that wild-type IRF3 bound to its cognate consensus sequence whereas IRF3-N85D failed to do so (Figure 4H). These results collectively show that deamidation impairs the ability of IRF3 to bind to its responsive element in promoters of inflammatory genes.

To determine the effect of IRF3 deamidation on SARS-CoV-2 replication, we first infected MEFs “reconstituted” with wild-type IRF3 and IRF3-N85D with GFP-marked VSV. We found that wild-type IRF3 markedly reduced VSV replication by immunofluorescence microscopy and plaque assay (Figure S4D and S4E). In contrast, IRF3-N85D only marginally reduced VSV replication. Next, we examined SARS-CoV-2 replication in these cells. To facilitate SARS-CoV-2 infection, we established MEF cell lines stably expressing human ACE2 (Figure S4F). These “reconstituted” MEFs were infected with SARS-CoV-2 and examined for innate immune response. Quantitative real-time analyses indicated that wild-type IRF3 induced modest level of expression of *Ifnb*, *Isg15*, *Isg56* and *Cxcl10* (Figure S4G). While IRF3-N85A robustly induced the expression of these genes, IRF3-N85D had minimal induction of these genes. Conversely, wild-type IRF3 and IRF3-N85A reduced viral RNAs by ~40% to 70%, while IRF3-N85D had no apparent effect on the RNA levels of *Nsp1* and *E*, or reduced *N* RNA by ~45% (Figure 4I). Plaque assay further showed that wild-type IRF3 and IRF3-N85A diminished infectious SARS-CoV in the medium by 75% and 90%, respectively, while IRF3-N85D reduced by ~30% (Figure 4J). Thus, deamidation at N85 impairs IRF3’s ability to defeat SARS-CoV-2 replication.

### SARS-CoV-2 Nsp8 and ORF8 Induce IRF3 Deamidation via CTPS1

To probe the virus-host interaction underpinning CTPS1-mediated IRF3 deamidation, we screened for viral proteins that interact with CTPS1 using a SARS-CoV-2 expression library. A co-IP assay identified multiple SARS-CoV-2 polypeptides that co-precipitated with CTPS1 in transfected 293T cells, including ORF7b, ORF8, M, Nsp8, Nsp10 and Nsp14 (Figure S5A). The interaction between CTPS1 and these SARS-CoV-2 proteins, except M, were further validated by co-IP assay using endogenous CTPS1 (Figure 5A). Three out of the six CTPS1-interacting SARS-CoV-2 polypeptides, i.e., ORF7b, ORF8 and Nsp8, also induced IRF3 deamidation in transfected 293T cells (Figure 1D). We reasoned that these SARS-CoV-2 polypeptides usurp CTPS1 to promote IRF3 deamidation. To test this, we depleted CTPS1 with shRNA and examined IRF3 deamidation. Indeed, depletion of CTPS1 shifted IRF3 toward the negative side of the gel strip (Figure 5B). Apparent shift of IRF3 was observed in the presence of ORF8 and NSP8, a minor portion of IRF3 was shifted when ORF7b was expressed when CTPS1 was depleted (Figure 5B). To further validate this, we compared wild-type IRF3 and the deamidation-resistant IRF3-N85A in two-dimension gel electrophoresis. This experiment revealed that wild-type IRF3, but not the IRF3-N85A, was shifted by ORF8 and Nsp8 expression (Figure 5C). Interestingly, ORF7b expression shifted both wild-type IRF3 and IRF3-N85A, suggesting that ORF7b likely induces the deamidation of IRF3 at sites other than N85. Nevertheless, these results show that ORF8 and Nsp8 induce the CTPS1-mediated deamidation of IRF3 at N85.

**Figure 5.**
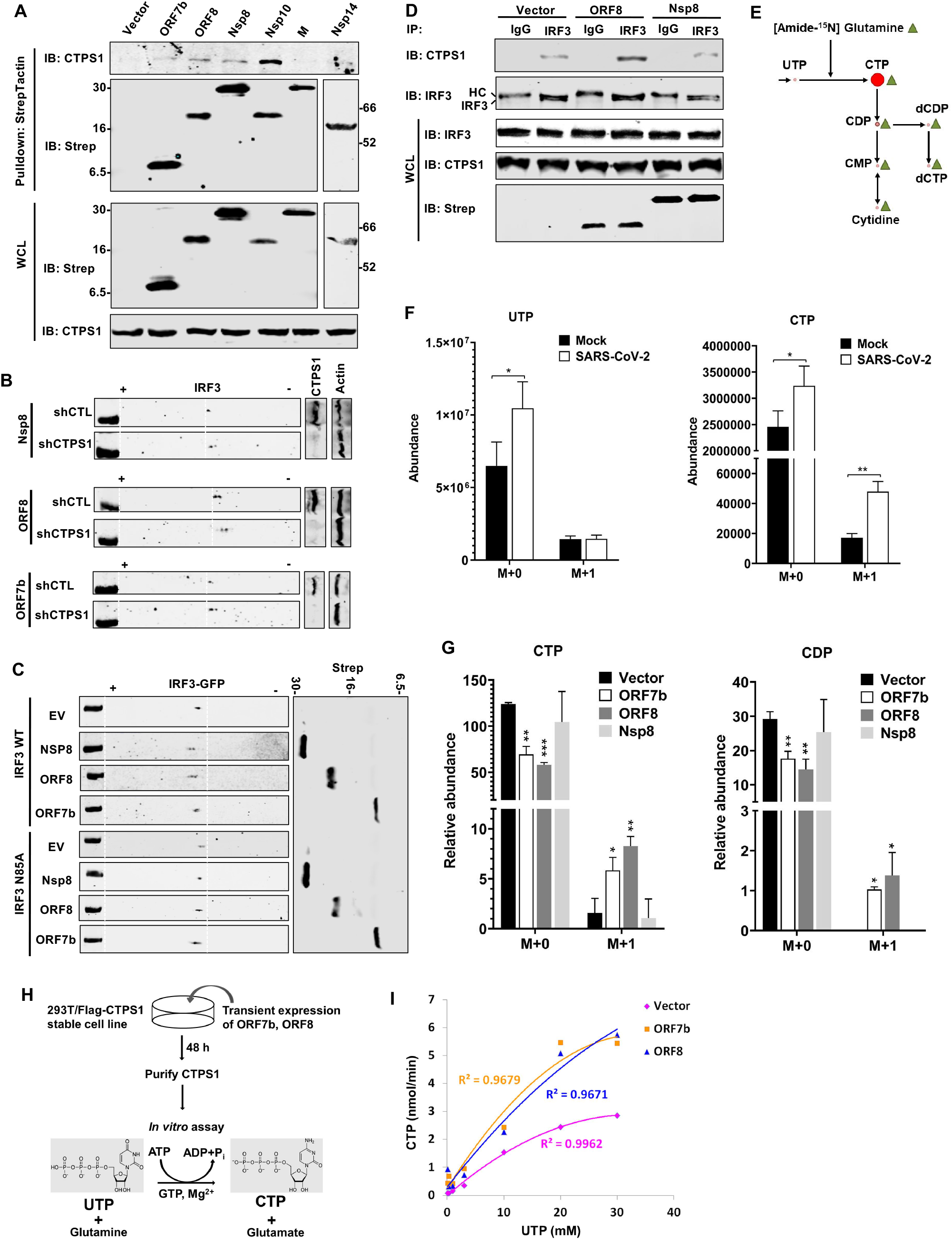
SARS-CoV-2 enhances CTPS1 enzymatic activity. (A) Interactions between endogenous CTPS1 and SARS-CoV-2 proteins were analyzed by co-immunoprecipitation in transfected 293T cells. Strep is a tag for SARS-CoV-2 proteins. (B) IRF3 charge in control (CTL) and CTPS1-depleted 293T cells, with the expression of SARS-CoV-2 proteins, was analyzed by two-dimensional gel electrophoresis and immunoblotting. (C) Effect of SARS-CoV-2 proteins on wild-type IRF3 and IRF3-N85A, in *IRF3*^*−/−*^ 293T cells expressing GFP-tagged IRF3-WT or IRF3-N85A, was determined by two-dimensional gel electrophoresis and immunoblotting. (D) Effect of SARS-CoV-2 ORF8 and Nsp8 on interaction between endogenous CTPS1 and IRF3, in 293T cells expressing ORF8 and Nsp8, was determined by co-immunoprecipitation and immunoblotting analyses. (E) Diagram of nitrogen incorporation in CTP synthesis using [Amide-^15^N]glutamine. (F) Intracellular UTP and CTP traced with [^15^N]glutamine were analyzed at 24 h after SARS-CoV-2 (MOI = 1) infection by mass spectrometry. M+a indicates the fraction of metabolites labeled with [^15^N]. M+2 is below the detection limit. (G) Effect of SARS-CoV-2 proteins on intracellular CTP and CDP traced with [^15^N]glutamine were determined by mass spectrometry in Caco-2 cells infected with lentivirus carrying SARS-CoV-2 ORF7b, ORF8, Nsp8 and control vector. Relative abundance of the metabolites was normalized by cell numbers. M+2 is below the detection limit. (H) Schematic diagram of the *in vitro* CTPS1 enzymatic assay. (I) Effect of SARS-CoV-2 ORF7b and ORF8 on kinetics of CTPS1 activity with respect to UTP, in the presence of 2 mM ATP, 2 mM L-glutamine, 0.1 mM GTP, was determined by *in vitro* enzymatic assay as described in (H) and analyzed by mass spectrometry. Error bars indicate SD of technical triplicates. Statistical significance was calculated using unpaired, two-tailed Student’s *t*-test. \**P* < 0.05; \*\**P* < 0.01; \*\*\**P* < 0.001. See related **Figure S5**.

To dissect the mechanism by which SARS-CoV-2 polypeptides promote CTPS1-mediated deamidation of IRF3, we determined whether ORF8 and Nsp8 impact the CTPS1-IRF3 interaction by co-IP assays. In 293T cells transiently expressing ORF8, more CTPS1 was precipitated by IRF3, indicating an elevated interaction between CTPS1 and IRF3 (Figure 5D). By contrast, Nsp8 had no effect on this interaction. Together, these results collectively show that SARS-CoV-2 polypeptides can enhance the ability of CTPS1 to deamidate IRF3.

### SARS-CoV-2 ORF8 and ORF7b Activate CTPS1 to Promote Nucleotide Synthesis

CTPS1 is responsible for the synthesis of CTP that is crucial for a balanced nucleotide pool during cell proliferation and viral replication. Activated nucleotide synthesis is likely to favor the transcription and genome replication of SARS-CoV-2. We then examined the metabolite of the glycolysis and nucleotide synthesis pathways. In the colorectal Caco-2 cell line that supports SARS-CoV-2 replication, we found that SARS-CoV-2 infection had no significant effect on the intracellular concentration of CTP (Figure S5B). However, the relative concentrations of UTP and UDP, and to a lesser extent UMP, immediate precursors of CTP, was significantly increased in Caco-2 cells at 72 h after SARS-CoV-2 infection (Figure S5C). Strikingly, CTP and UTP were significantly decreased at 96 h post-infection. These results support the rate-limiting role of CTP synthetases in catalyzing UTP to CTP conversion and suggest the decrease of CTP and UTP at 96 hpi is likely due to a rapid consumption. To determine the rate of synthesis that reflects the activity of CTPS1, we analyzed CTP synthesis using isotope tracing with [^15^N]glutamine that donates [^15^N]amide to CTP (Figure 5E). Compared to mock-infected cells, SARS-CoV-2 increased the labeled (M+1) CTP by >2-fold in Caco-2 cells with a 30-minute tracing (Figure 5F). Interestingly, SARS-CoV-2 infection had no apparent effect on the [^15^N]UTP (M+1) under similar conditions, indicating the specificity of CTPS1 activation during SARS-CoV-2 infection. [^15^N]UTP (M+1) is the product of the *de novo* pyrimidine synthesis where CAD catalyzes dihydroorotate synthesis using glutamine. Next, we established Caco-2 cell lines that stably express SARS-CoV-2 polypeptides, including ORF7b, ORF8 and Nsp8. When flux analysis with [^15^N]glutamine was performed, we found that cells expressing ORF7b and ORF8 had >3- and 5-fold more [^15^N]CTP (M+1) compared to control cells (Vector group), respectively (Figure 5G). Consistently, ORF7b and ORF8 also increased [^15^N]CDP (M+1), an immediate product hydrolyzed from CTP. Strikingly, Nsp8 expression had no apparent effect on labeled [^15^N]CTP. Robust increase in [^15^N]CTP was also observed in ORF8-expressing LoVo colorectal cells (Figure S5E). These results show that ORF7b and ORF8 promote CTP synthesis.

To probe the effect of ORF7b and ORF8 on the enzymatic activity of CTPS1, we purified CTPS1 from stable 293T/CTPS1 cells with transient expression of ORF7b and ORF8, and performed biochemical assays to determine the kinetic parameters kcat and Km of CTPS1 (Figure 5H). Compared to the control group (vector), ORF7b and ORF8 expression increased k_cat_ of CTPS1 by ~ 1-fold and ~ 2-fold, respectively (Figure 5I and S5F). ORF8, but not ORF7b, increased Km of CTPS1 by ~ 0.7 fold. These results demonstrate that SARS-CoV-2 ORF7b and ORF8 activate CTPS1 to synthesize CTP.

### Inhibitors of CTPS1 Impede SARS-CoV-2 Replication

Based on the roles of CTPS1 elucidated above, inhibition of CTPS1 is expected to diminish CTP supply and restore IFN induction, thereby impeding SARS-CoV-2 replication. We thus sought to develop small-molecule inhibitors to further verify CTPS1’s role and furnish therapeutic candidates to combat SARS-CoV-2 infection and COVID-19. Because the GAT domain within CTPS1 has cysteine hydrolase activity and contains a catalytic cysteine (C399) in its active site, CTPS1 is sensitive to covalent inhibition by electrophilic compounds exemplified by DON (Goto et al., 2004; Huang et al., 2017). We thus screened a panel of electrophilic analogues to search for inhibitors of CTPS1. Our screening effort led to the identification of compound **1** as an effective CTPS1 inhibitor (Figure 6A). As analyzed by two-dimensional gel electrophoresis, compound **1** demonstrated efficacy in inhibiting IRF3 deamidation in Caco-2 cells expressing ORF8, or A549 and LoVo cells expressing Nsp8 (Figure 6B and S6A). Accordingly, compound **1** increased *IFNB1* expression in a dose-dependent manner in 293T cells infected with Sendai virus (Figure S6B). To determine the specificity of compound **1**, we depleted CTPS1 in 293T cells for a luciferase report assay. We found that compound **1** elevated IFN induction in control 293T cells, but failed to do so in CTPS1-depleted 293T cells and had no effect on NF-κB activation (Figure 6C and S6C). To validate that compound **1** targets CTPS1, we employed compound **2**, a close relative of compound **1** containing a terminal alkyne tag, for biochemical labeling using 293T cells expressing CTPS1 (Figure 6A). This assay showed that compound **2** covalently labeled CTPS1 in a dose-dependent manner (Figure 6D). Taken together, these results demonstrate that compound **1** inhibits CTPS1 to elevate IFN induction.

**Figure 6.**
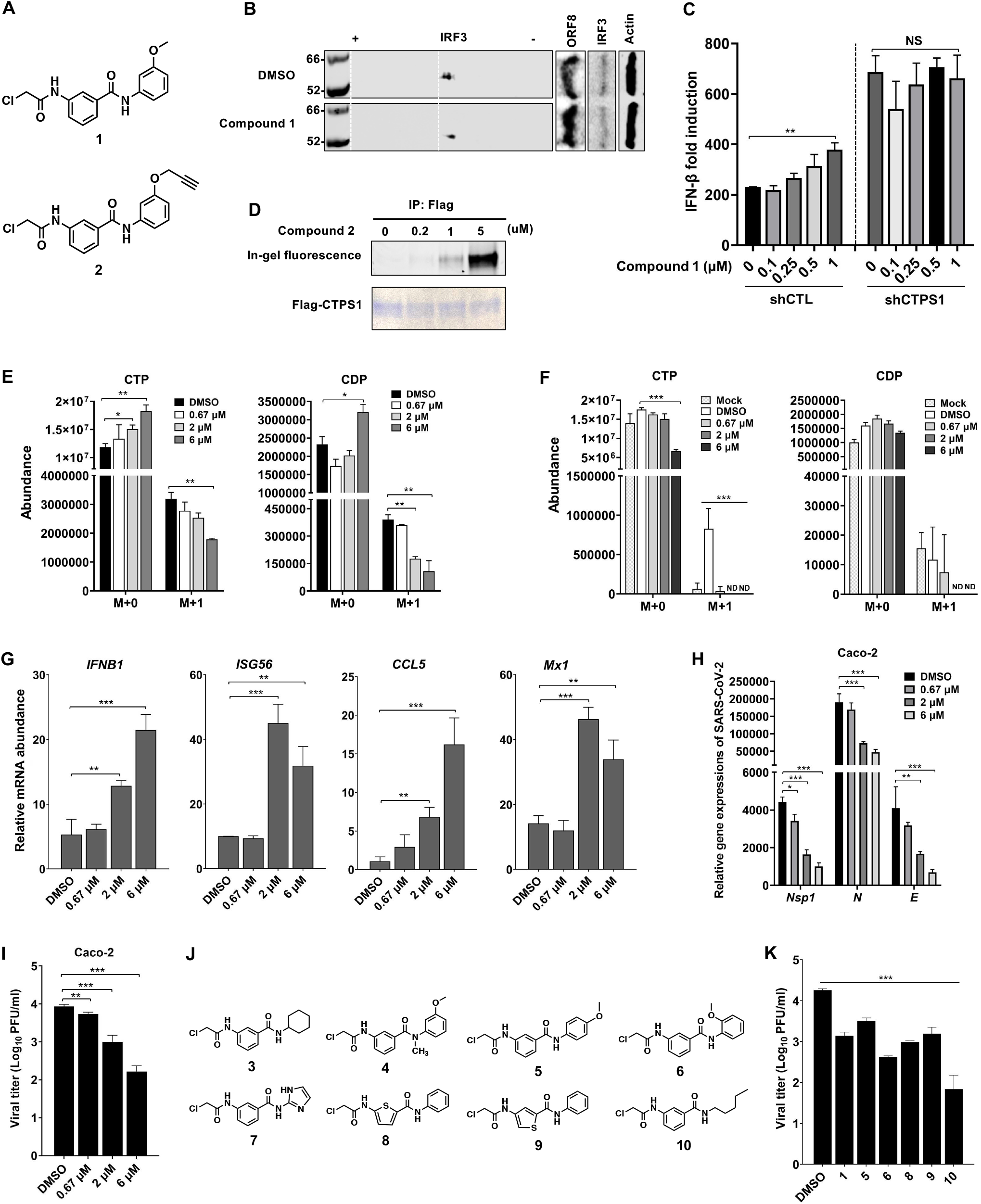
CTPS1 inhibitors suppress SARS-CoV-2 replication. (A) Structures of Compound **1** and **2**. (B) Effect of Compound **1** on SARS-CoV-2 ORF8-induced IRF3 deamidation was analyzed by two-dimensional gel electrophoresis and immunoblotting in ORF8-expressing Caco-2 cells with Compound **1** (5 μM) treatment. (C) Effect of Compound **1** on IFN induction by Sendai virus infection was determined by luciferase reporter assay using control (CTL) or CTPS1-depleted 293T cells treated with increasing amount of Compound **1**. (D) Flag-CTPS1 expressed 293T cells were treated with Compound 2 (5 μM) for 2 h. CTPS1 was purified and subjected to binding analysis by in-gel fluorescence imaging and Coomassie blue staining. (E and F) Effect of Compound **1** on intracellular CTP and CDP traced with [^15^N]glutamine was determined by mass spectrometry using SARS-CoV-2 ORF8-expressing Caco-2 cells (E) or SARS-CoV-2-infected Caco-2 cells (F) treated with increasing amounts of compound **1**. M+2 was below the detection limit. (G - I) Caco-2 cells were treated with Compound **1** and infected with SARS-CoV-2 (MOI = 0.1). The mRNA abundance of antiviral genes was determined by real-time PCR at 48 h after SARS-CoV-2 infection (G). Effect of Compound **1** on SARS-CoV-2 RNA abundance (H) and infectious viral progeny (I) was determined at 72 h after SARS-CoV-2 infection by real-time PCR analysis of total RNA and plaque assay of the medium, respectively. (J) Structures of Compound **1** derivatives. (K) Effect of Compound **1** and its derivatives on SARS-CoV-2 replication was determined by plaque assay at 72 h post-infection (MOI = 0.1) in medium of Caco-2 cells. Error bars indicate SD of technical triplicates. Statistical significance was calculated using unpaired, two-tailed Student’s *t*-test. \**P* < 0.05; \*\**P* < 0.01; \*\*\**P* < 0.001. See related **Figure S6**.

Activated CTPS1 also increases CTP supply in cells infected with SARS-CoV-2 to facilitate viral replication. With Caco-2 cells that stably express ORF8, we performed [^15^N]glutamine flux analysis with compound **1** treatment. As shown in Figure 6E, compound **1** treatment diminished [^15^N]CTP, and to a much greater extent [^15^N]CDP, in a dose-dependent manner. Similar reductions were observed in ORF8-expressing LoVo cells after compound **1** treatment (Figure S6D). Remarkably, compound **1** potently diminished the intracellular concentration of [^15^N]CTP and [^15^N]CDP in Caco-2 cells infected with SARS-CoV-2 (Figure 6F). These results show that an inhibitor of CTPS1 can block CTP synthesis in SARS-CoV-2-infected cells and in cells expressing ORF8.

To probe the biological consequence of compound **1** treatment, we analyzed the expression of antiviral genes, including *IFNB1*, *ISG56*, *CCL5* and *Mx1*, in SARS-CoV-2-infected Caco-2 cells. Real-time PCR analysis indicated that compound **1** at the concentrations of 2 μM and 6 μM elevated the expression of these antiviral genes (Figure 6G). Consistent with the elevated antiviral gene expression, the abundance of viral RNAs, including *Nsp1*, *N* and *E*, was reduced by compound **1** in a dose-dependent manner (Figure 6H), with more than 50% and 75% reduction at the concentrations of 2 μM and 6 μM, respectively. The reduced viral RNA abundance also correlated with lower viral yield, in which compound **1** reduced viral yield by ~10- and 100-fold at the concentrations of 2 μM and 6 μM (Figure 6I). Similar results were observed in SARS-CoV-2-infected NHBE cells when treated with compound **1**, including elevated antiviral gene expression and reduced viral RNA and yield (Figure S6E-S6G). These results collectively show that compound **1** inhibits CTPS1 to impede nucleotide synthesis and restore IFN induction, with synergistical effects that diminish SARS-CoV-2 replication.

To improve the antiviral potency of compound **1**, we designed and synthesized eight structural analogues, compound 3 – 10 (Figure 6J). NMR and mass spectra of compound **3**-**10** are consistent with their chemical structures. Cell toxicity test showed that these molecules had no significant effects on cell viability up to 3 μM and reduced cell viability and proliferation at 9 μM, likely due to the diminished CTP supply when CTPS1 was inhibited (Figure S6H). An IFN induction reporter assay showed that four of the eight derivatives, including compound **5**, **6**, **9** and **10**, had improved effect to increase IFN induction, compared with compound **1** (Figure S6I). Whereas compound **3** and **4** had no effect on IFN induction, compound **7** and **8** demonstrated modest effect. Thus, we selected five derivatives, including compound **5**, **6**, **8**, **9** and **10**, in this regard for further SARS-CoV-2 study. Consistent with their ability to enhance IFN induction, compound **6** and **10** showed more robust antiviral activity in SARS-CoV-2 infection than compound **1**, as determined by plaque assay (Figure 6K). The result of plaque assay also correlated with SARS-CoV-2 RNA abundance in Caco-2 cells (Figure S6J). These results identified a number of CTPS1 inhibitors that potently antagonize SARS-CoV-2 replication in cultured cells. The efficacy of these molecules in diminishing SARS-CoV-2 replication is being examined using rodent models.

## DISCUSSION

SARS-CoV-2 has caused a global pandemic with a record of more than 100 million infections, 2.16 million deaths and an unknown number of asymptomatic cases. Studies involving cultured cells, model animals and COVID-19 patients indicate that SARS-CoV-2 effectively inhibits the production of type I and III interferons (Johansen et al., 2020; Park and Iwasaki, 2020; Sa Ribero et al., 2020). However, the molecular mechanism by which SARS-CoV-2 does so is not well understood. Earlier works comparing SARS-CoV-2 genome sequences to those of other beta coronaviruses, particularly SARS-CoV and MERS-CoV, predict putative viral polypeptides in modulating host innate immune defense, including IFN induction (Lei et al., 2020; Li et al., 2020). Here, we report that SARS-CoV-2 deploys multiple proteins to activate CTPS1, which promotes CTP synthesis, while inactivating IRF3 and muting IFN induction. Remarkably, pharmacological inhibition of CTPS1 potently impedes CTP synthesis and effectively restores IFN induction, thereby diminishing SARS-CoV-2 replication and offering an antiviral strategy targeting a host enzyme.

Dysregulated immune response is a characteristic shared among COVID-19 patients under severe and critical conditions (Coperchini et al., 2020; Tang et al., 2020). Among the skewed cytokine profile, type I IFNs are produced at very low or under detection levels in most severe or critical COVID-19 patients, which likely contributes to the rapid replication of SARS-CoV-2 in these patients. Surprisingly, recent reports indicate that the majority of severe and critical COVID-19 patients show low inflammatory cytokine levels, compared to patients infected with influenza virus (Mudd et al., 2020). To dissect the mechanism of innate immune evasion by SARS-CoV-2, we first showed that RNA produced from SARS-CoV-2-infected NHBE cells potently induced IFN, whereas SARS-CoV-2 failed to do so during infection, unlike Sendai virus, suggesting that SARS-CoV-2 viral proteins antagonize IFN induction. Indeed, a screen utilizing a SARS-CoV-2 expression library identified ORF7b, ORF8, Nsp8 and Nsp13 as inhibitors of IFN induction. Further analysis showed that these SARS-CoV-2 polypeptides target IRF3 for post-translational modification. Given the ubiquitous role of type I IFNs in host innate immune defense against viral infection, regulatory mechanisms governing IRF3 activation are expected to operate independent of cell type and tissue origin (Ivashkiv and Donlin, 2014; Stetson and Medzhitov, 2006). Upstream components, such as pattern recognition receptors and their cognate adaptors, may be tissue- and cell type-specific (Amarante-Mendes et al., 2018; Mogensen, 2009). Accordingly, viral factors targeting these upstream components likely function in a tissue-dependent manner, and those meddling with the downstream components, such as IRF3, are anticipated to work independent of tissues or organs. In the lung, the epithelial cells and pneumocytes of the airway and respiratory track are the first responders in IFN production during SARS-CoV-2 infection (Sa Ribero et al., 2020; Zamorano Cuervo and Grandvaux, 2020).

The SARS-CoV-2 polypeptides that induce IRF3 deamidation are relatively small and unlikely to function as intrinsic deamidases. Indeed, a focused shRNA-mediated screen targeting cellular glutamine amidotransferases (GATs) identified CTPS1 as a negative regulator of IFN induction. CTPS1 belongs to the cellular GAT family that is known for metabolic functions in biosynthesis of cellular building blocks in preparation for proliferation (Choi and Carman, 2007; F. Massière and Badet-Denisot., 1998). CTPS1 demonstrates intrinsic activity to deamidate IRF3 *in vitro* and in cells, which is dependent on the active site required for the glutamine-hydrolysis (glutaminase) activity in catalyzing CTP synthesis. This study adds CTPS1 to the growing list of protein deamidases that were originally known as cellular GATs, expanding the functional repertoire of protein deamidation and GATs in immune regulation (He et al., 2015; Li et al., 2019; Zhang et al., 2018; Zhao et al., 2016b). Interestingly, CTPS1 and CTPS2 share 74% amino acid homology and were predicted to be functionally redundant (Kassel et al., 2010). However, we found that CTPS1, but not CTPS2, interacts with IRF3 in cells, suggesting that these two closely related enzymes are functionally distinct. Indeed, loss or deficiency of CTPS1 due to mutations was found to impair CTP synthesis in T cell proliferation and result in primary immune deficiency, despite the fact that CTPS2 is highly expressed in T cells (Martin et al., 2014). Given the pivotal roles of CTPS1 in T cell-mediated adaptive immunity, it remains an interesting question whether the protein-deamidating activity of CTPS1 is important for T cell immune function.

Deamidation results in the loss of DNA-binding activity of IRF3 to its cognate sequences, supporting the role of deamidation in diminishing IFN induction by IRF3. This constitutes a strategy by which viruses effectively shut down antiviral gene expression during infection. Similarly, cells may deploy this mechanism to curtail expression of genes that are not essential during proliferation when CTPS1 is highly active. Such a mechanism is analogous to the CAD-mediated RelA deamidation that shunts RelA to transactivate the expression of key glycolytic enzymes in promoting carbon metabolism during S phase (Zhao et al., 2020). SARS-CoV-2 hijacks CTPS1 to deamidate IRF3 to evade IFN induction during infection, which may explain previous observations that SARS-CoV-2 fails to induce IFN production in COVID-19 patients and in animal models (Johansen et al., 2020). Intriguingly, deamidated IRF3, similar to deamidated RelA, still translocates into the nucleus, suggesting that deamidated IRF3 may have unidentified functions relevant to biological processes in the nucleus.

Nucleotide supply is a rate-limiting factor for cell proliferation and virus replication (Mayer et al., 2019; Zhu and Thompson, 2019). As intracellular obligate pathogens, viruses rely on cellular machinery for their macromolecular biosynthesis (Eisenreich et al., 2019; Mesquita I and J., 2018). During viral productive infection, nucleotides are used for transcription, translation (ribosome regeneration), genome replication and lipid synthesis for the assembly and maturation of virion progeny. Not surprisingly, viruses often activate metabolic enzymes to fuel nucleotide synthesis in support of their replication (Sanchez and Lagunoff, 2015). We discovered that SARS-CoV-2 infection and the expression of ORF7b and ORF8 activate CTPS1 to promote the *de novo* CTP synthesis, thereby fueling viral replication. Strikingly, activated CTPS1 also inhibits type I IFN induction via deamidating IRF3. Thus, SARS-CoV-2 couples the inhibition of type I IFN induction to CTP synthesis via activating CTPS1. This predicts that SARS-CoV-2 relies on CTPS1 for its replication, and conversely inhibiting CTPS1 likely impedes SARS-CoV-2 replication. Indeed, we found that depletion and pharmacological inhibition of CTPS1 greatly diminished CTP synthesis and effectively restored antiviral IFN induction in SARS-CoV-2 infection.

Given the scarcity of antiviral therapeutics against SARS-CoV-2 and COVID-19, we developed small-molecule inhibitors of CTPS1 as antiviral candidates. These molecules stimulate IFN induction, but not NF-κB activation. CTPS1 and CAD negatively regulate the IFN and NF-κB induction, respectively (Zhao et al., 2020). The specific stimulation of IFN induction by compound **1** and its derivative thus suggests their inhibition of CTPS1, but not CAD. Furthermore, the effect of compound **1** on IFN induction was observed in wild-type cells and this effect was abolished in CTPS1-depleted cells, supporting the conclusion that compound 1 inhibits CTPS1 to boost IFN production. Indeed, a derivative of compound 1 carrying a reactive warhead cross-links with CTPS1 in vitro. These results collectively show that compound 1 targets CTPS1 to promote IFN induction. Given that CTPS1 is essential for cell proliferation, compound 1 potentially induces toxicity in proliferating cells. The premise is that SARS-CoV-2 activates CTPS1 to facilitate its replication, which permits the selective inhibition of CTPS1 with low dose of compound 1 or one of its derivatives. Additionally, we cannot exclude the possibility that compound 1 and its derivatives target cellular proteins other than CTPS1. Moreover, application of CTPS1 inhibitors has to be optimized to avoid toxicity to host cells, T cells for adaptive immunity. Future experiments will be necessary to optimize the conditions that minimize the side effects of these CTPS1 inhibitors using animal models.

## ACKNOWLEDGMENTS

We thank Dr. Zhiwei Liao for RNA-sequencing analysis, Drs. Jae-Jung and Woo-Jin Shin for cell lines. We thank Dr. Lucio Comai and Mrs. Jill Henley for the assistance on experiments using BSL3 facility. This work is partly supported by grants from NIH (DE027556, CA221521 and DE026003), startup funds from the Herman Ostrow School of Dentistry of USC, USC Zumberge Epidemic and Virus Award and National Natural Science Foundation of China (32070678 to Taijiao Jiang). Bianca Espinosa was supported by the National Science Foundation Graduate Research Fellowship Program (DGE 1418060).

## STAR★ METHODS

Key Resources Table.

Key Resources Table
| REAGENT or RESOURCE | SOURCE | IDENTIFIER |
| --- | --- | --- |
| **Antibodies** |  |  |
| Mouse monoclonal Flag M2 antibody | Sigma | Cat#: F3165 |
| mouse monoclonal β-Actin antibody | Abcam | Cat#: ab8226 |
| Rabbit polyclonal IRF3 antibody | Santa Cruz | Cat#: sc-33641 |
| phospho-IRF3 (Ser396) antibody | Cell Signaling Technology | Cat#: 4D4G |
| phospho-IRF3 (Ser386) antibody | Abcam | Cat#: Ab76493 |
| Rabbit polyclonal CTPS1 antibody | Proteintech | Cat#: 15914-1-AP |
| Mouse Strep monoclonal antibody | BioLegend | Cat#: 688202 |
| Rabbit polyclonal GST antibody | Santa Cruz | Cat# sc-459 |
| **Chemicals** |  |  |
| [α-P<sup>32</sup>]-ATP | PerkinElmer | Cat#: BLU003H250UC |
| [<sup>15</sup>N] Glutamine | Cambridge Isotope Lab | Cat#: NLM-557-1 |
| ATP | Thermo Fisher Scientific | Cat#: R0441 |
| GTP | Thermo Fisher Scientific | Cat#: R0461 |
| UTP | Thermo Fisher Scientific | Cat#: J63427 |
| StrepTactin agarose | GE Healthcare | Cat#: 28-9355-99 |
| 3-Aminophenol | Sigma-Aldrich | Cat#: 100242 |
| Propargyl bromide | Sigma-Aldrich | Cat#: P51001 |
| Chloroacetyl chloride | Sigma-Aldrich | Cat#: 104493 |
| *m*-Anisidine | Sigma-Aldrich | Cat#: A88204 |
| 3-Nitrobenzoic acid | Fisher Scientific | Cat#: AC128451000 |
| Cyclohexylamine | TCI | Cat#: C0494 |
| *p*-Anisidine | Sigma-Aldrich | Cat#: A88255 |
| *o*-Anisidine | Sigma-Aldrich | Cat#: A88182 |
| 2-Aminoimidazole sulfate | Sigma-Aldrich | Cat#: 197912 |
| Thiophene-2-carboxylic acid | AmBeed | Cat#: A103725 |
| amylamine | TCI | Cat#: A0445 |
| PEG-2-biotin-azide | Conju-Probe | Cat#: CP-3040 |
| **Critical Commercial Assays** |  |  |
| Human IFN-β ELISA kit | PBL assay science | Cat#: 41410 |
| Human CCL5 ELISA kit | R&D systems | Cat#: DRN00B |
| Mouse IFN-β ELISA kit | PBL Assay Science | Cat#: 42400 |
| Mouse CXCL10 ELISA kit | R&D systems | Cat#: DY466-05 |
| **Experimental Models: Cell lines** |  |  |
| HEK293T | ATCC | Cat#: ACS-4500 |
| THP1 | ATCC | Cat#: TIB-202 |
| A549 | Laboratory of Jae Jung | N/A |
| LoVo | Laboratory of Chengyu Liang | N/A |
| Caco-2 | ATCC | Cat#: HTB-37 |
| Calu-3 | ATCC | Cat#: HTB-55 |
| NHBE | ATCC | Cat#: PCS-300-010 |
| IRF3/7<sup>-/-</sup> MEF | Laboratory of P. Feng | N/A |
| Vero E6 | Laboratory of Jae Jung | N/A |
| **Viruses** |  |  |
| Sendai virus (SeV) | Charles River | PI-1, SV |
| Vesicular stomatitis virus (VSV) | Laboratory of P. Feng | N/A |
| SARS-CoV-2 | Laboratory of Jae Jung | N/A |
| Oligonucleotides |  |  |
| qPCR primers, see Table S1 | N/A | N/A |
| ChIP-qPCR primers, see Table S1 | N/A | N/A |
| sgRNA primers, see Table S1 | N/A | N/A |
| Human GATs shRNA | Thermo fisher Scientific | (Li et al., 2019;<br>Zhao et al., 2020) |

## EXPERIMENTAL MODEL AND SUBJECT DETAILS

### Cell Culture

HEK293T, A549, LoVo, *Irf3*^*−/−*^*Irf7*^*−/−*^ mouse embryonic fibroblasts (MEFs) were cultured in Dulbecco’s modified Eagle’s medium (DMEM, Hyclone). THP1 cells were cultured in RPMI 1640 medium. Caco-2, Calu-3 were cultured in MEM medium. All these cell lines were supplemented with 10% fetal bovine serum (FBS, HyClone), penicillin (100 U/mL) and streptomycin (100 μg/mL), and maintained at 37 ℃ in a humidified atmosphere of 5% CO_2_. Primary normal, human bronchial/tracheal epithelial cells (NHBE) were cultured in airway epithelial cell medium according to the ATCC recommendation.

### Viruses

Sendai virus (SeV) was purchased from Charles River. Vesicular stomatitis virus (VSV) was amplified using Vero cells. SARS-CoV-2 was propagated in Vero E6 cells. All SARS-CoV-2 related viral propagation, viral infection, and vital titration were performed in biosafety level 3 (BSL-3) facility (USC).

SARS-CoV-2 propagation: Vero E6 cells were seeded at 1.5 × 10^6^ cells per T25 flask for 12 h. Cells were washed with FBS-free DMEM medium once, and infected with SARS-CoV-2 at MOI 0.005 in FBS-free DMEM medium. Cells were checked daily for cytopathic effect (CPE). Virus-containing medium was harvested when virus-induced CPE reached approximately 80% (around 72 h after viral infection). Centrifuge 3000 rpm, 5 min, and store at −80 ℃.

SARS-CoV-2 infection: NHBE (1.5 × 10^5^ cells), Calu-3 (5 × 10^5^ cells) or Caco-2 cells (2 × 10^5^ cells) were seeded in one well of 12-well plates. Cells were washed with FBS-free medium before viral infection. SARS-CoV-2 were diluted in 250 μl (per well) medium corresponding to the cell line. Viral infection was incubated on a rocker for 45 min at 37°C. Cells were washed with fresh medium, and medium containing 10% FBS was added.

SARS-CoV-2 viral titration (plaque assay): Vero E6 cells were seeded in 6- or 12-well plates. When cell confluence reaches to 100%, cells were washed with FBS-free medium, and infected with serially diluted SARS-CoV-2. After infection, medium was removed, and overlay medium containing FBS-free 1 × DMEM and 1% low-melting point agarose was added. At 72 h post infection, cells were fixed with 4% paraformaldehyde (PFA) overnight, and stained with 0.2% crystal violet. Plaques were counted on a light box.

### Plasmids

Luciferase reporter plasmids for IFN-β, NF-κB promoters, RIG-I-N, MAVS, TBK1, IRF3-5D and shRNA for human glutamine amidotransferases (CTPS1, CTPS2, GFPT1, GFPT2, GMPS, PFAS, PPAT, CPS1, ASNS and NADSYN1) were described previously (Li et al., 2019; Zhao et al., 2020; Zhao et al., 2016b). A cDNA construct was used to amplify and clone CTPS1 into mammalian expression vectors. Point mutants of IRF3 and CTPS1, including IRF3-Q15E, IRF3-N85D, IRF3-N184D, IRF3-N217D, IRF3-N389D, IRF3-N397D, IRF3-N85A, IRF3-N85Q, and CTPS1 enzyme-deficient (CTPS1-ED) mutant (C399A/H526A/E528A) were generated by site-directed mutagenesis and confirmed by sequencing. Lentiviral expression constructs containing IRF3, CTPS1 and hACE2 were generated from pCDH-CMV-EF1-Puro or pCDH-CMV-EF1-Hygro by molecular cloning. pLVX-EF1alpha-2XStrep-IRES-Puro containing SARS-CoV-2 viral genes described previously and provided by Dr. Nevan J. Krogan (Gordon et al., 2020).

## METHODS DETAILS

### Quantitative Real-time PCR (qRT-PCR)

qRT-PCR was performed as previously described (Zhao et al., 2020). Briefly, total RNA was extracted from mock- or virus-infected cells using TRIzol reagent (Invitrogen). cDNA was synthesized from one microgram total RNA using reverse transcriptase (Invitrogen) according to the manufacturer’s instruction. Quantitative real-time PCR (qRT-PCR) reaction was performed with SYBR Green Master Mix (Sigma) or qPCRBIO SyGreen Blue Mix Lo-ROX (Genesee Scientific). Gene expression level was calculated by 2^−ΔΔCt^ method. Primers for qRT-PCR were listed in Table S1.

### Lentivirus-mediated Stable Cell Line Construction

Lentivirus production was carried out in HEK293T cells. Briefly, 293T cells were co-transfected with packaging plasmids (VSV-G, DR8.9) and pCDH lentiviral expression vector or lentiviral shRNA plasmids. At 48 h post transfection, the medium was harvested and filtered. HEK293T, *Irf3*^*−/−*^*Irf7*^*−/−*^ mouse embryonic fibroblasts (MEFs), Caco-2, LoVo and A549 cells were infected with the supernatant, supplied with polybrene (8 ug/ml), and centrifuged at 1800 rpm for 50 min at 30℃. Cells were incubated at 37℃ for 6 h, and maintained in DMEM with 10% FBS. Selection was performed at 48 h post infection with puromycin (1-2 μg/ml) or hygromycin (200 μg/ml).

To establish IRF3 knockout cell line, 293T cells were transduced with lentivirus expressing sgRNA targeting IRF3 (pL-CRISPR.EFS.PAC-Targeting-IRF3, Table S1) and selected with 1 μg/ml puromycin. Single colonies were isolated and screened by immunoblotting with IRF3 antibody.

### Dual-Luciferase Reporter Assay

HEK293T cells in 24-well plates (~50% cell density) were transfected with reporter plasmid cocktail containing 50 ng luciferase reporter plasmids (ISRE-luc, IFN-β-luc or NF-κB), 5 ng TK-renilla luciferase reporter (control vector) and the indicated expression plasmids by calcium phosphate precipitation. Whole cell lysates were prepared at 24 – 30 h post-transfection, and used for dual luciferase assay according to the manufacturer’s instruction (Promega).

### Confocal Microscopy Analysis

*Irf3*^*−/−*^*Irf7*^*−/−*^ MEFs reconstituted with Flag-IRF3 wild-type, Flag-IRF3-N85D were infected with or without SeV (100 HA units/ml). Sixteen hours later, cells were washed, fixed as previous report (Rao et al., 2015). Celle were incubated with primary mouse monoclonal anti-Flag antibody and Alex Fluor 488 congugated goat secondary antibody, and analyzed with confocal microscope (Leica).

### RNA Sequencing

*Irf3*^*−/−*^*Irf7*^*−/−*^ MEFs reconstituted with Flag-IRF3 wild-type, Flag-IRF3-N85D were mock-infected or infected with SeV (100 HA units/ml). Total RNA was extracted at 12 h post-infection. RNA sequencing was performed using Hiseq3000 at the Technology Center for Genomic & Bioinformatics of UCLA. RNA sequencing results were analyzed as previously described in our publication (Zhao et al., 2020).

### Protein Expression and Purification

HEK293T cells were transfected with Flag-tagged or GST-Tagged genes of interest. Cells were harvested at 48 h post transfection, and lysed with Triton X-100 buffer (20 mM Tris, pH 7.5, 150 mM NaCl, 1 mM EDTA, 20 mM β-glycerophosphate, 10% glycerol) supplemented with a protease inhibitor cocktail (Roche). Whole cell lysates (WCLs) were sonicated, incubated at 4℃ for 30 min on a rotator, and centrifuged at 12,000 rpm for 30 min. Supernatant was filtered, precleared with sepharose 4B agarose beads (Thermo) at 4℃ for 1 h. The pre-cleared WCLs were incubated with anti-FLAG M2 agarose beads or glutathione-conjugated agarose beads at 4℃ for 4 h. Anti-FLAG M2 magnetic beads were washed extensively with lysis buffer and eluted with 0.2 mg/ml 3xFlag peptide. GST beads were extensively washed and used immediately for *in vitro* on-column deamidation assay. Concentration of purified proteins was analyzed by SDS-PAGE and Coomassie staining, with BSA as a standard.

### Two-dimensional Gel Electrophoresis

Cells (1 × 10^6^) were resuspended in 150 μl rehydration buffer (8 M Urea, 2% CHAPS, 0.5% IPG buffer, 0.002% bromophenol blue), sonicated three times, and incubated 15 min on ice. Whole cell lysates were centrifuged at 12,000 g for 15 min. Supernatants were loaded to IEF strips for isoelectric focusing with a program comprising: 20 V, 10 h (rehydration); 500 V, 1 h; 1000 V, 1 h; 1000-5000 V, 4 h; 5000 V, 4 h. Then, strips were incubated with SDS equilibration buffer (50 mM Tris-HCl [pH8.8], 6 M urea, 30% glycerol, 2% SDS, 0.001% Bromophenol Blue) containing 10 mg/ml DTT for 15 min and SDS equilibration buffer containing 2-iodoacetamide for 15 min. Strips were washed with SDS-PAGE buffer, resolved by SDS-PAGE, and analyzed by immunoblotting.

### In vitro Deamidation Assay

Expression plasmids containing IRF3-WT-GST, IRF3-N85D-GST, Flag-CTPS1 were transfected into HEK293T cells. Cell lysates were prepared at 48 h post transfection and proteins were purified with anti-FLAG M2 agarose (Sigma) and glutathione-conjugated agarose beads (Sigma). *In vitro* on-column deamidation of IRF3 was performed as previously reported (Zhao et al., 2020). Briefly, 0.2 μg of CTPS1 and 0.6 μg of IRF3-WT-GST or IRF3-N85D-GST (on beads) were added to a total volume of 50 μl. The reaction was carried out at 37 ℃ for 45 min in deamidation buffer (50 mM Tris-HCl at pH 8.0, 20 mM MgCl_2_, 5 mM KCl, 1 mM ATP, 1 mM GTP). IRF3-WT-GST or IRF3-N85D-GST were eluted with rehydration buffer (8 M Urea, 2% CHAPS, 0.5% IPG Buffer, 0.002% bromophenol blue) at room temperature. Samples were analyzed by two-dimensional gel electrophoresis and immunoblotting.

### Mass Spectrometry Analysis for Deamidation Sites

To identify deamidation sites, HEK293T cells were transfected with plasmid containing IRF3-GST without or with that containing the enzyme-deficient CTPS1 mutant (Flag-CTPS1-ED). Transfected cells were harvested at 48 h post transfected and IRF3-GST was purified with glutathione-conjugated agarose beads from whole cell lysates. Purified proteins were subjected to SDS-PAGE and Coomassie blue staining. Gel slices containing IRF3-GST were prepared for in-gel digestion and mass spectrometry analysis (Poochon Scientific).

### Metabolic Profiling and Isotope Tracing

Caco-2 cells were mock-infected or infected with SARS-CoV-2 at MOI = 1. Cells were harvested at 6 h, 24 h, 48 h and 72 h post-infection for metabolomics analysis.

Isotope tracing experiments were performed as previously described (Zhao et al., 2020). To analyze the effect of SARS-CoV-2 proteins on nucleotide synthesis, LoVo and Caco-2 cell lines stably expressing SARS-CoV-2 ORF7b, ORF8 and Nsp8 were cultured with medium containing [^15^N-amide]glutamine for 30 min and 1 h. Cells were washed with 1 ml ice-cold ammonium acetate (NH_4_AcO, 150 mM, pH 7.3), added 1 ml −80℃ cold MeOH, and incubated at −80 ℃ for 20 min. After incubation, cells were scraped off and supernatants were transferred into microfuge tubes. Samples were pelleted at 4 ℃ for 5 min at 15k rpm. The supernatant was transferred into new microfuge tubes, dried at room temperature under vacuum, and re-suspended in water for LC-MS run.

Samples were randomized and analyzed on a Q-Exactive Plus hybrid quadrupole-Orbitrap mass spectrometer coupled to Vanquish UHPLC system (Thermo Fisher). The mass spectrometer was run in polarity switching mode (+3.00 kV/-2.25 kV) with an m/z window ranging from 65 to 975. Mobile phase A was 5 mM NH_4_AcO, pH 9.9, and mobile phase B was acetonitrile. Metabolites were separated on a Luna 3 μm NH2 100 NH2 100A° (150 × 2.0 mm) column (Phenomenex). The flow rate was 0.3 ml/min, and the gradient was from 15% A to 95% A in 18 min, followed by an isocratic step for 9 min and re-equilibration for 7 min. All samples were run in biological triplicate. Metabolites were detected and quantified as area under the curve based on retention time and accurate mass (5 ppm) using the TraceFinder 4.1 (Thermo Scientific) software. Raw data was corrected for naturally occurring ^15^N abundance.

### CTPS1 Enzymatic Activity Assay

Flag-CTPS1 expressed 293T stable cell line was transfected with plasmids containing SARS-CoV-2 ORF7b, ORF8, Nsp8 and empty vector for 40 h. CTPS1 was purified with anti-FLAG M2 agarose via one-step affinity chromatography. CTPS1 activity was determined by measuring the conversion of UTP to CTP via mass spectrometry. The standard reaction mixture containing 50 mM Tris-HCl (pH 8.0), 10 mM MgCl2, 10 mM 2-mercaptoethanol, 2 mM L-glutamine, 1 mM GTP, 1 mM ATP, increasing amount of UTP (0 to 30 mM), and an appropriate dilution of CTPS1 in a total volume of 50 μl. The reactions were equilibrated to 37 ℃ for 45 min, and quenched by adding 250 μl cold (−80 ℃) methanol and incubating at −80 ℃ for 20 min. The metabolites were analyzed by LS-MS as described above.

### Enzyme-linked Immunosorbent Assay (ELISA)

Control and CTPS1-depleted 293T or THP1 cells, *Irf3*^*−/−*^*Irf7*^*−/−*^ MEF reconstituted with IRF3-WT, IRF3-N85D, IRF3-N85A and empty vector were infected with SeV (100 HAU/ml). Medium of cells was harvested at the indicated time points. Human IFN-β, CCL5, and mouse IFN-β, CXCL10 were analyzed by commercial ELISA kits according to the manufacturer’s instruction.

### Electrophoresis Mobility Shift Assay (EMSA)

Flag-IRF3-WT and Flag-IRF3-N85D were purified from HEK293T cells. DNA-binding was performed as previously described (Andrilenas et al., 2018). Binding reactions were carried out in 20 μl volumes containing: 2.5 nM P32 labeled DNA probe targeting IFN-β promoter (forward: GCACCGCTAACCGAAACCGAAACTGTGC; reverse: GCACAGTTTCGGTTTCGGTTAGCGGTGC); 10 mM Tris pH 7.5; 50 mM NaCl; 2 mM DTT and indicated purified Flag-IRF3-WT or Flag-IRF3-N85D. Unlabeled probe (cold probe) was used for competition assay. Reactions were incubated at room temperature for 45 min, and resolved in 6% polyacrylamide gels (29:1 crosslinking) with 0.5 × TBE running buffer at 200 V/cm on ice until the loading dye front reached the bottom of the gel. Gels were dried and analyzed using phosphorimaging instrumentation.

### Small Molecule Synthesis

Reagents and solvents were obtained from commercial suppliers and used without further purification, unless otherwise stated. Flash column chromatography was carried out using an automated system (Teledyne Isco CombiFlash. Reverse phase high performance liquid chromatography (RP-HPLC) was carried out on a Shimadzu HPLC system. All anhydrous reactions were carried out under nitrogen atmosphere. NMR spectra were obtained on Varian VNMRS-500, VNMRS-600, or Mercury-400.

#### Synthesis of compounds 1-10

##### Step 1

**X** (1 eq.) and **Y** (1.2 eq.) were dissolved in DCM/DMF (4:1). HBTU (4 eq.) and triethylamine (4 eq.) were added and the solution was allowed to stir at room temperature for 16 hours. After 16 hours, the reaction mixture was diluted with EtOAc and washed with 10% Na_2_CO_3_ followed by brine 3 times. The organic layer was dried with Na_2_SO_4_ and concentrated by rotary evaporation. The residue was purified via flash column chromatography (EtOAc/Hex). All products moved to **Step *2*** except for the **compound 4** intermediate, which first moved to **Step 1.2**.

**Table 1.** Synthesis of target compounds

| Compound | X | Y |
| --- | --- | --- |
| 1 | 3-nitrobenzoic acid | m-anisidine |
| 2 | 3-nitrobenzoic acid | 3-(prop-2ynyloxy)aniline |
| 3 | 3-nitrobenzoic acid | Cyclohexamine |
| 4 | 3-nitrobenzoic acid | m-anisidine |
| 5 | 3-nitrobenzoic acid | p-anisidine |
| 6 | 3-nitrobenzoic acid | o-anisidine |
| 7 | 3-nitrobenzoic acid | bis(1h-imidazol-2-amine;sulfuric acid |
| 8 | 5-nitrothiophene-2-carboxylic acid | Aniline |
| 9 | 4-nitrothiophene-2-carboxylic acid | Aniline |
| 10 | 3-nitrobenzoic acid | Amylamine |

###### Step 1.2 (*only applies to compound 4*)

The intermediate for **compound 4** from **Step 1** (1 eq) was dissolved in dry THF and cooled to 0°C. NaH (60% in oil) (1.5 eq) was added portion wise to the stirring solution. Methyl iodide (1.1 eq.) was added drop wise. The reaction mixture was allowed to warm to room temperature. The round bottom was transferred to oil bath and refluxed at 75°C for 2 hrs. The reaction mixture was poured in ice water and was extracted with EtOAc 3 times. The organic layer was washed with brine 2 times, dried with Na_2_SO_4_, and concentrated by rotary evaporation. The residue was purified via flash column chromatography and the intermediate moved onto **Step 2**.

##### Step 2

Intermediates from **Steps 1 and 1.2** (1 eq.) were dissolved in MeOH. Zn (5 eq.) and NH_4_Cl (5 eq.) were added, and the mixture was allowed to stir at room temperature for 16 hr. The reaction mixture was dissolved in EtOA and washed with 10% Na_2_CO_3_ followed by brine 3 times. The organic layer was dried over NaSO_4_ and concentrated by rotary evaporation to yield the intermediates which were used in the next reaction without further purification.

##### Step 3

Intermediates from **Step 3** (1 eq.) were dissolved in anhydrous DCM/THF (1:4) under nitrogen gas. DIPEA (1.2 eq) was added via syringe. Chloroacetyl chloride (1.2 eq) was added via syringe slowly dropwise. The reaction mixture was allowed to stir overnight. The mixture was diluted with EtOAc and washed with 10% Na_2_CO_3_. The organic layer was dried over NaSO_4_ and the residue was purified via flash chromatography (EtOAc/Hexane) to yield the final products.

### Drug Treatment

For SARS-CoV-2 infection, Caco-2 or NHBE cells were pre-treated with compound 1 or its derivatives (5, 6, 8, 9, and 10) for 2 h. Then the medium were removed. Cells were washed and infected with SARS-CoV-2. Afterwards, medium containing virus was removed, and cells were cultured with drug-containing medium. Drugs were added at each 24 h after viral infection until the end of the experiments. DMSO was used as control. To test the effect of compound 1 on IRF3 deamidation, Caco-2 cells expressing ORF8, or A549 and LoVo cells expressing Nsp8 were treated with 5 μM compound 1 for 4 h. To analyze the effect of compound 1 on intracellular metabolites, ORF8 expressed Caco-2 or LoVo cells were treated with 5 μM compound 1 for 2 h.

