## Supplemental Figures and legends for "Targeting CTP Synthetase 1 to Restore Interferon Induction and Impede Nucleotide Synthesis in SARS-CoV-2 Infection"

**Affiliations:**
[1]Section of Infection and Immunity, Herman Ostrow School of Dentistry, Norris Comprehensive Cancer Center, University of Southern California, Los Angeles, CA 90089, USA;
[2]Department of Chemistry, Dornsife College of Arts, Letters and Sciences, University of Southern California, Los Angeles, CA 90089, USA;
[3]Florida Research and Innovation Center, Cleveland Clinic, FL 34987, USA;
[4]Mork Family Department of Chemical Engineering and Materials Science, Norris Comprehensive Cancer Center, Los Angeles, CA 90089, USA;
[5]Center for Systems Medicine, Institute of Basic Medical Sciences, Chinese Academy of Medical Sciences & Peking Union Medical College, Beijing 100005; Suzhou Institute of Systems Medicine, Suzhou, Jiangsu 215123; Bioland Laboratory (Guangzhou Regenerative Medicine and Health Guangdong Laboratory), 510005 Guangzhou, China.
[6]Lead Contact.

[a]Current address: Department of Surgery, Cedars-Sinai Medical Center, Los Angeles, CA 90048.

### Figure S1

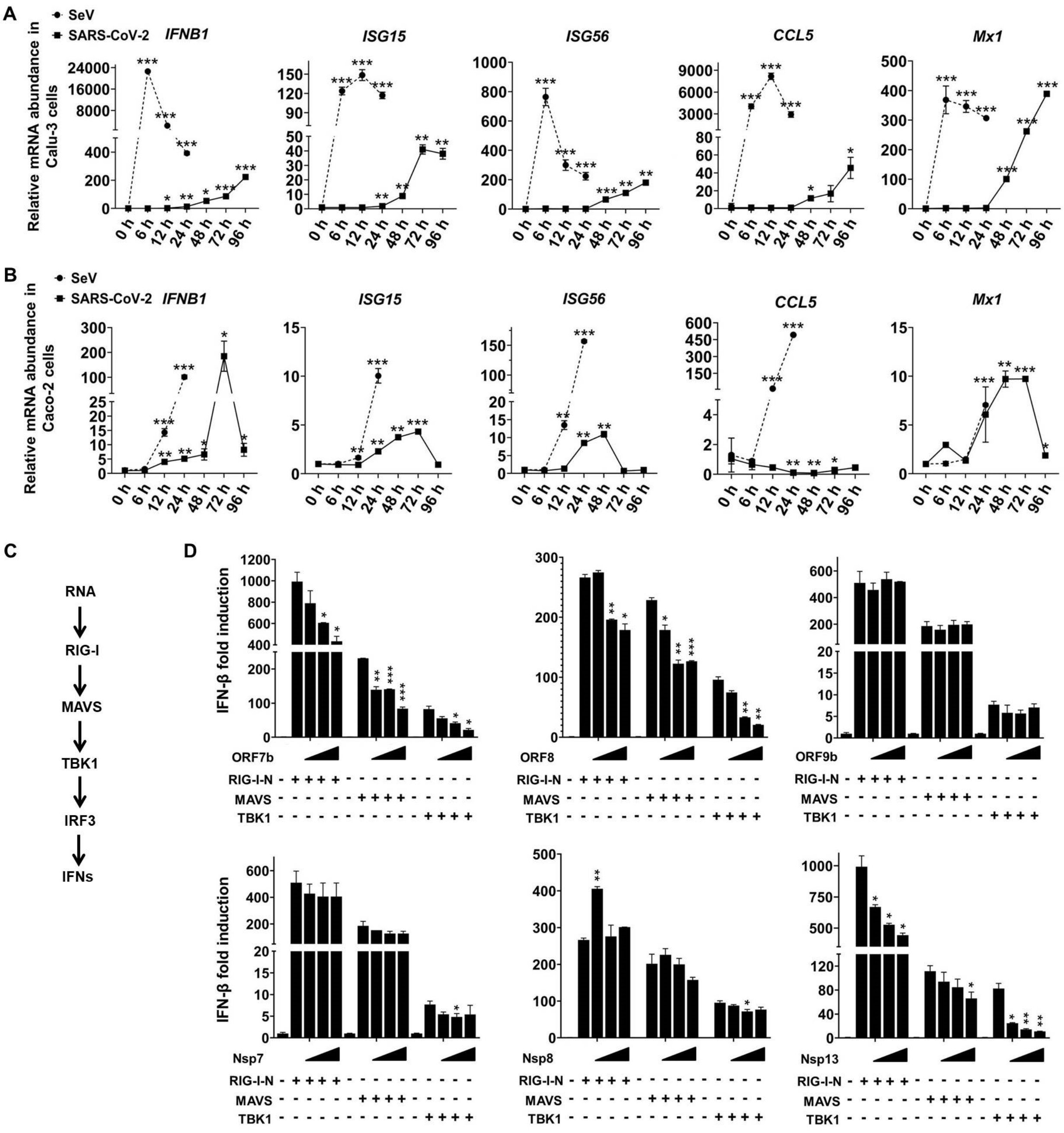
A
SeV
SARS-CoV-2
IFNB1
ISG15
ISG56
CCL5
Mx1
Relative mRNA abundance in Calu-3 cells
0 h 6 h 12 h 24 h 48 h 72 h 96 h
B
Relative mRNA abundance in Caco-2 cells
C
RNA
RIG-I
MAVS
TBK1
IRF3
IFNs
D
IFN-β fold induction
ORF7b
ORF8
ORF9b
Nsp7
Nsp8
Nsp13
RIG-I-N
MAVS
TBK1

**Figure S1. SARS-CoV-2 inhibits IFN induction,** Related to **Figure 1**

(A and B) Calu-3 and Caco-2 cells were infected with Sendai virus (SeV) (100 HAU/ml) or SARS-CoV-2 (MOI = 1). Total RNA was extracted, reverse transcribed and analyzed by real-time PCR with primers specific for *IFNB1*, *ISG15*, *ISG56*, *CCL5* and *Mx1*.

(C) Diagram of the RIG-I-IFN pathway that can be triggered by RNA virus infection.

(D) Effect of SARS-CoV-2 proteins on IFN-β induction was determined by luciferase reporter assay in 293T cells transfected with IFN-β reporter cocktail, plasmids containing increasing doses of SARS-CoV-2 viral proteins and indicated components of the RIG-I-IFN pathway.

Error bars indicate SD of technical triplicates. Statistical significance was calculated using unpaired, two-tailed Student's *t*-test. $^*P < 0.05$; $^{**}P < 0.01$; $^{***}P < 0.001$.

### Figure S2

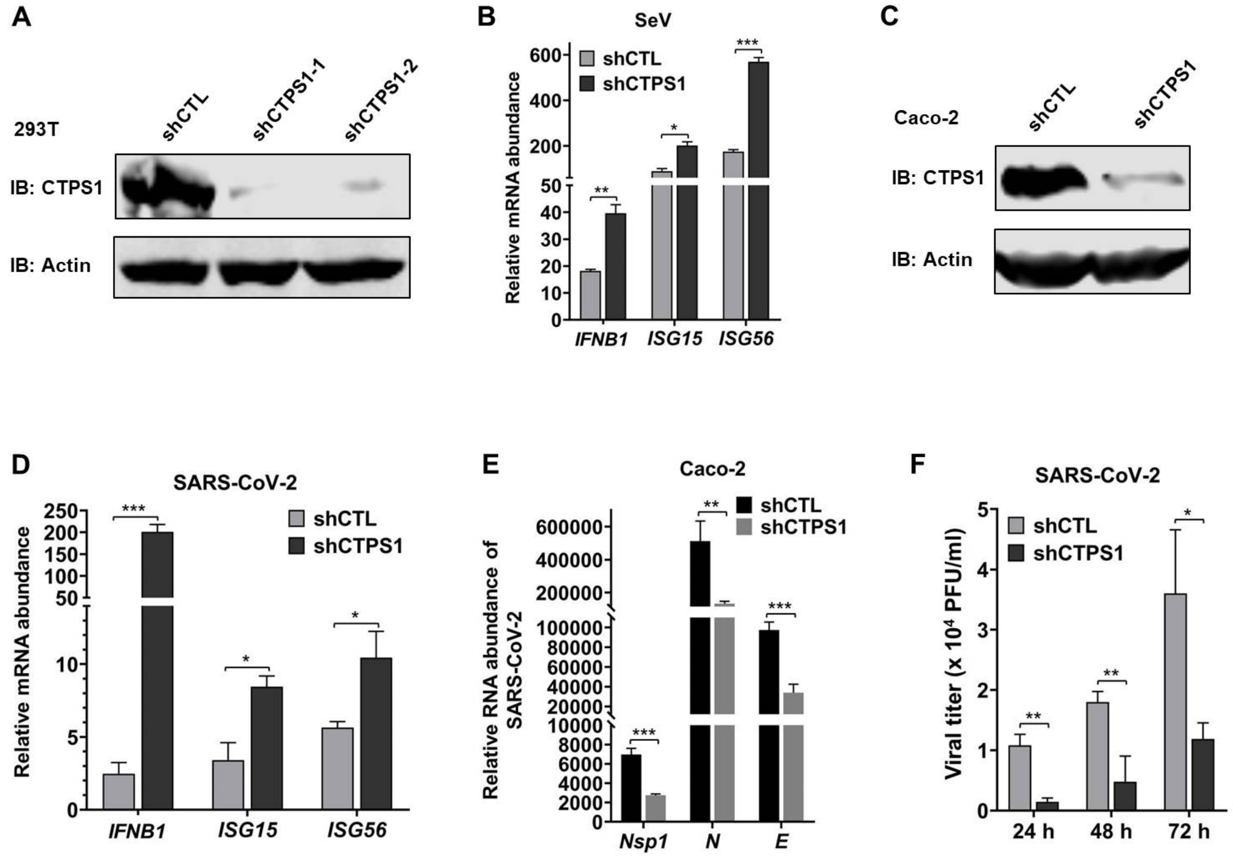

A
293T
shCTL
shCTPS1-1
shCTPS1-2
IB: CTPS1
IB: Actin
B
SeV
shCTL
shCTPS1
Relative mRNA abundance
IFNB1
ISG15
ISG56
C
Caco-2
shCTL
shCTPS1
IB: CTPS1
IB: Actin
D
SARS-CoV-2
shCTL
shCTPS1
Relative mRNA abundance
IFNB1
ISG15
ISG56
E
Caco-2
shCTL
shCTPS1
Relative RNA abundance of SARS-CoV-2
Nsp1
N
E
F
SARS-CoV-2
shCTL
shCTPS1
Viral titer (x 10⁴ PFU/ml)
24 h
48 h
72 h

**Figure S2. CTPS1 inhibits IFN induction.** Related to **Figure 2**

(A) Knockdown of CTPS1 in 293T cells was determined by immunoblotting using cells infected with lentivirus containing control (CTL) or CTPS1 shRNA.

(B) The mRNA abundance of antiviral genes induced by Sendai virus infection was determined by real-time PCR at 12 h post-infection using CTPS1 depleted and control (CTL) THP1 cells.

(C) Knockdown of CTPS1 in Caco-2 cells was determined by immunoblotting using cells infected with lentivirus containing control (CTL) or CTPS1 shRNA.

(D) Effect of CTPS1 depletion on the expression of cellular antiviral genes was determined by real-time PCR at 24 h post-SARS-CoV-2 (MOI = 0.5) infection in CTPS1-depleted and control Caco-2 cells.

(E and F) CTPS1-depleted and control Caco-2 cells were infected with SARS-CoV-2 (MOI 0.1) for 72 h. Effect of CTPS1 depletion on viral gene expression was determined by real-time PCR analysis of total RNA. Viral titer in the medium was measured by plaque assay.

Error bars indicate SD of technical triplicates. Statistical significance was calculated using unpaired, two-tailed Student's *t*-test. *$P < 0.05$; **$P < 0.01$; ***$P < 0.001$.

### Figure S3

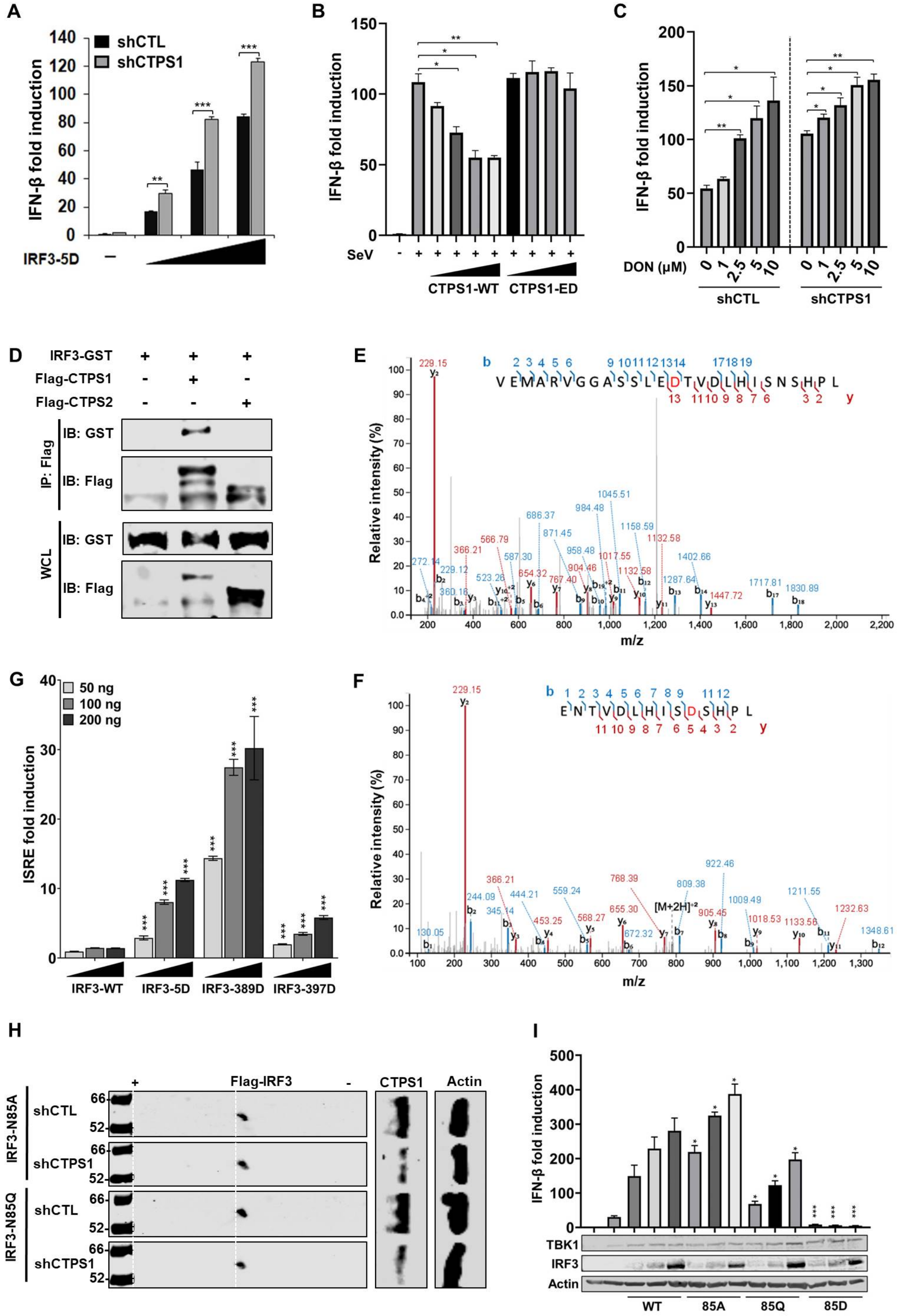

A
IFN-β fold induction
shCTL
shCTPS1
IRF3-5D
B
IFN-β fold induction
SeV
CTPS1-WT
CTPS1-ED
C
IFN-β fold induction
DON (μM)
shCTL
shCTPS1
D
IRF3-GST
Flag-CTPS1
Flag-CTPS2
IP: Flag
IB: GST
IB: Flag
WCL
E
Relative intensity (%)
m/z
VEMARVGGASSLEDTVDLHISNSHPL
F
Relative intensity (%)
m/z
ENTVDLHISDSHPL
G
ISRE fold induction
50 ng
100 ng
200 ng
IRF3-WT
IRF3-5D
IRF3-389D
IRF3-397D
H
Flag-IRF3
CTPS1
Actin
IRF3-N85A
IRF3-N85Q
shCTL
shCTPS1
I
IFN-β fold induction
TBK1
IRF3
Actin
WT
85A
85Q
85D

**Figure S3. CTPS1 deamidates IRF3.** Related to **Figure 3**

(A) IFN-β promoter activity was determined by reporter assay in 293T cells transfected with IFN-β reporter plasmid cocktail and expression plasmids containing increasing amounts of IRF3-5D.

(B) Modulation of IFN-β induction was determined by promoter activity in 293T cells expressing CTPS1-WT and CTPS1-ED, with Sendai virus infection.

(C) Effect of 6-diazo-5-oxo-L-norleucine (DON) on IFN-β induction was determined by luciferase reporter assay in CTPS1-depleted and control (CTL) 293T cells, with Sendai virus infection.

(D) Interactions between IRF3 and CTPS1 or CTPS2 were analyzed by co-immunoprecipitation and immunoblotting in 293T cells transfected with plasmids containing GST-IRF3, and Flag-CTPS1 or Flag-CTPS2.

(E and F) IRF3 deamidation was determined by tandem mass spectrometry using affinity purified IRF3 in the presence of CTPS1-ED. The m/z spectrums of the peptide containing N389D (E) and N397D (F) are shown with the deamidated D residue highlighted in red.

(G) Effects of several deamidations on IRF3 were determined by luciferase assay using 293T cells transfected with the ISRE reporter cocktail and plasmids containing increasing amounts of wild-type IRF3 or indicated deamidated IRF3 mutants.

(H) IFN-β promoter activity was determined in 293T cells transfected with IFN-β reporter plasmid cocktail and expression plasmids containing TBK1, and increasing amounts of IRF3-WT and mutants. Expression levels of the components were analyzed by immunoblotting.

(I) Effect of CTPS1 on IRF3-N85A and IRF3-N85Q was determined by two-dimensional gel electrophoresis and immunoblotting with control (CTL) and CTPS1-depleted 293T cells transfected with a plasmid containing IRF3-N85A or IRF3-N85Q.

Error bars indicate SD of technical triplicates. Statistical significance was calculated using unpaired, two-tailed Student's *t*-test. $^*P < 0.05$; $^{**}P < 0.01$; $^{***}P < 0.001$.

### Figure S4

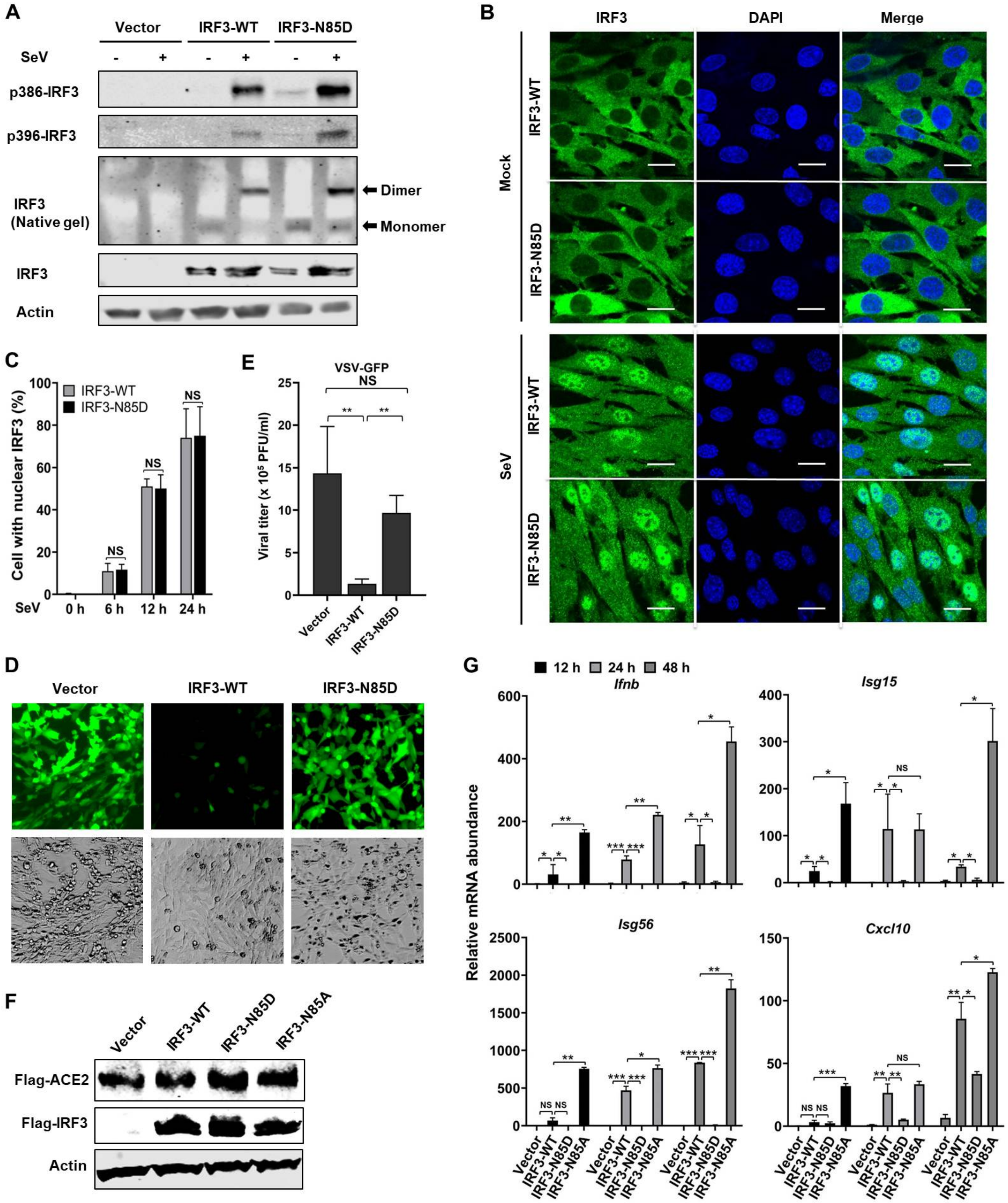
A
Vector
IRF3-WT
IRF3-N85D
SeV
p386-IRF3
p396-IRF3
IRF3
(Native gel)
Dimer
Monomer
IRF3
Actin
B
IRF3
DAPI
Merge
Mock
SeV
IRF3-WT
IRF3-N85D
C
Cell with nuclear IRF3 (%)
IRF3-WT
IRF3-N85D
NS
SeV
0 h
6 h
12 h
24 h
E
VSV-GFP
Viral titer (x 10⁵ PFU/ml)
Vector
IRF3-WT
IRF3-N85D
D
Vector
IRF3-WT
IRF3-N85D
F
Vector
IRF3-WT
IRF3-N85D
IRF3-N85A
Flag-ACE2
Flag-IRF3
Actin
G
12 h
24 h
48 h
Relative mRNA abundance
Ifnb
Isg15
Isg56
Cxcl10
Vector
IRF3-WT
IRF3-N85D
IRF3-N85A

**Figure S4. Deamidation impedes IRF3 to activate antiviral immune responses by blocking its DNA binding activity.** Related to **Figure 4**

(A) IRF3-WT and IRF3-N85D reconstituted *Irf3*$^{-/-}$*Irf7*$^{-/-}$ MEF cells were infected with SeV for 12 h. Phosphorylation and dimerization of IRF3 were resolved by SDS-PAGE or native PAGE with the whole cell lysates, and followed by immunoblotting analysis with indicated antibodies.

(B and C) IRF3-WT and IRF3-N85D reconstituted *Irf3*$^{-/-}$*Irf7*$^{-/-}$ MEF cells were infected with or without Sendai virus. Subcellular localization of IRF3-WT and IRF3-N85D were analyzed at 24 h post-infection by immunofluorescence using a confocal microscope. Scale bars, 5 μm (B). Cells with IRF3-WT or IRF3-N85D nuclear localization were counted at indicated time points after Sendai infection. Error bars indicate SD. NS, no significance (C).

(D and E) IRF3-WT and IRF3-N85D reconstituted *Irf3*$^{-/-}$*Irf7*$^{-/-}$ MEF cells were infected with VSV-GFP (MOI 0.01) for 10 h. GFP-positive cells were recorded with fluorescence microscopy (D). Viral replication in medium was determined by plaque assay (E).

(F) *Irf3*$^{-/-}$*Irf7*$^{-/-}$ MEF cells were infected with lentivirus containing Flag-tagged human ACE2, selected with hygromycin, then reconstituted with Vector, IRF3-WT, IRF3-N85D or IRF3-N85A. Expression levels of ACE2 and IRF3 were analyzed by immunoblotting.

(G) RF3-WT, IRF3-N85D, IRF3-N85A and vector reconstituted *Irf3*$^{-/-}$*Irf7*$^{-/-}$ MEF cells expressing human ACE2 were infected with SARS-CoV-2 (MOI = 0.1). The mRNA abundance of *Ifnb*, *Isg15*, *Isg56* and *Cxcl10* were analyzed by real-time PCR at indicated time points post-infection.

Error bars indicate SD of technical triplicates. Statistical significance was calculated using unpaired, two-tailed Student's *t*-test. NS, no significance; $^{*}P < 0.05$; $^{**}P < 0.01$; $^{***}P < 0.001$.

### Figure S5

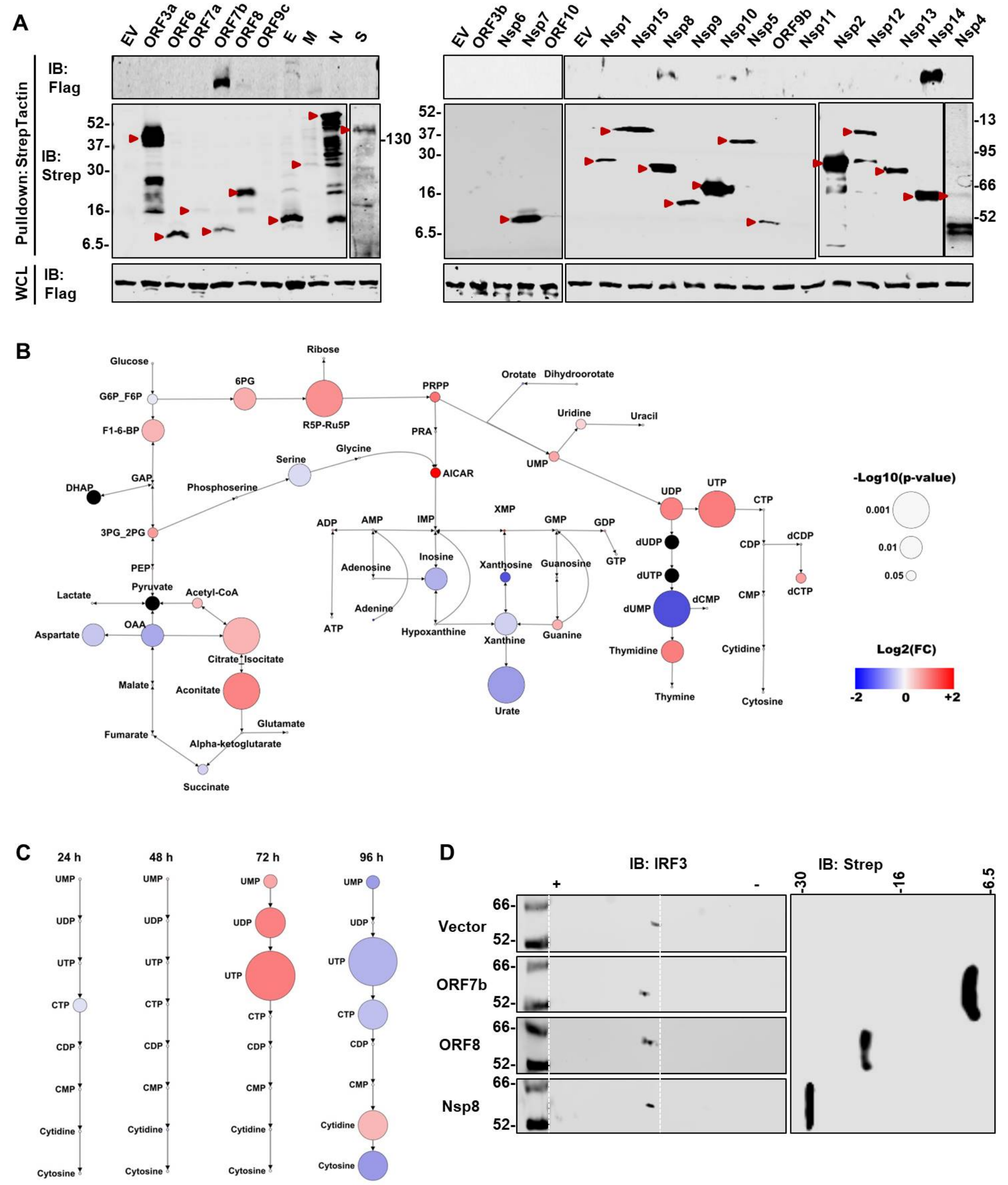


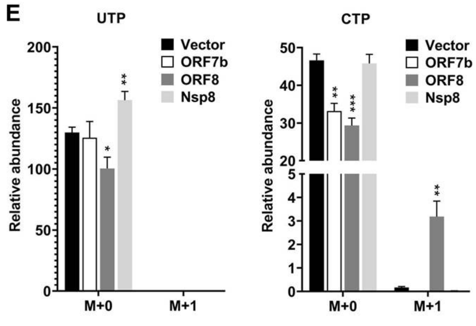


F

| | $K_m$ (mM) | $K_{cat}$ ($min^{-1}$) |
|---|---|---|
| Vector | 30.94 ± 10.45 | 0.44 ± 0.09 |
| ORF7b | 30.54 ± 21.53 | 0.88 ± 0.36 |
| ORF8 | 55.20 ± 50.29 | 1.27 ± 0.82 |

**Figure S5. SARS-CoV-2 promotes CTP synthesis by targeting CTPS1.** Related to **Figure 5**

(A) Interaction between CTPS1 and SARS-CoV-2 proteins were determined by co-immunoprecipitation and immunoblotting in 293T cells transfected with plasmid containing Flag-CTPS1 and indicated strep-tagged SARS-CoV-2 proteins. Strep indicates SARS-CoV-2 proteins.

(B and C) Caco-2 cells were infected with SARS-CoV-2 with MOI = 1 for 24 h, 48 h, 72 h and 96 h. Global metabolite profile was analyzed by mass spectrometry. SARS-CoV-2-induced metabolite changes in the central carbon pathway at 72 h post-infection are shown as a metabolic map (B). SARS-CoV-2-induced changes of pyrimidine synthesis at different time points are shown in (C).

(D) Effects of the selected SARS-CoV-2 proteins on IRF3 deamidation were analyzed by two-dimensional gel electrophoresis and immunoblotting using Caco-2 cells infected with lentivirus carrying SARS-CoV-2 ORF7b, ORF8, Nsp8 or control vector.

(E) Effects of SARS-CoV-2 proteins on intracellular UTP and CTP traced with [$^{15}$N]glutamine were determined by mass spectrometry in LoVo cells infected with lentivirus carrying SARS-CoV-2 ORF7b, ORF8, Nsp8 or control vector. Relative abundance of the metabolites was normalized by cell numbers. M+2 is below the detection limit.

(F) Kinetic constants for CTP synthetase, related to Figure 5I. $K_m$ and $K_{cat}$ were calculated by Michaelis-Menten equation.

Error bars indicate SD of technical triplicates. Statistical significance was calculated using unpaired, two-tailed Student's *t*-test. **P* < 0.05; ***P* < 0.01; ****P* < 0.001.

### Figure S6

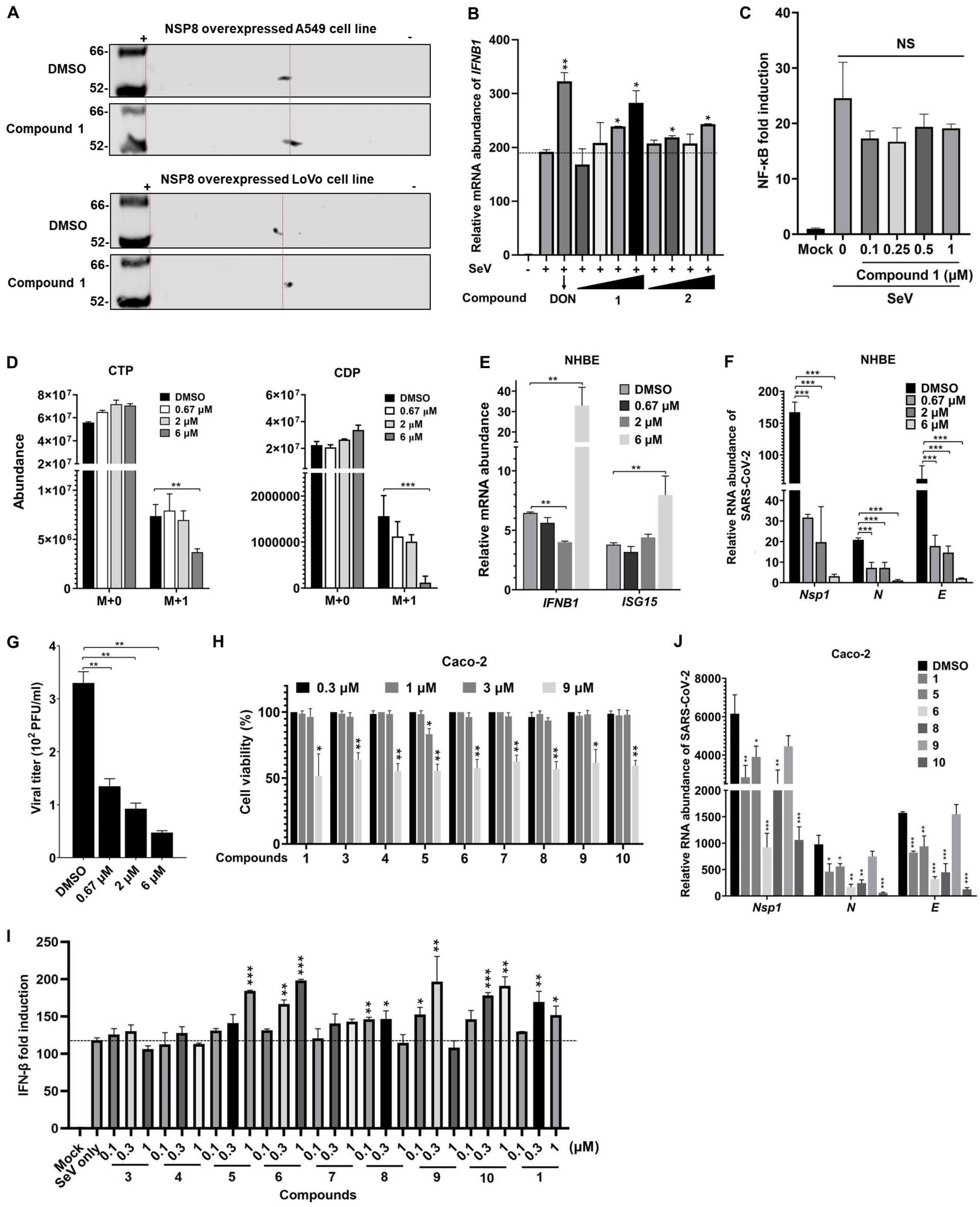

A
NSP8 overexpressed A549 cell line
NSP8 overexpressed LoVo cell line
DMSO
Compound 1
B
Relative mRNA abundance of IFNB1
SeV
Compound
DON
C
NF-κB fold induction
NS
Mock
Compound 1 (μM)
SeV
D
CTP
CDP
Abundance
DMSO
0.67 μM
2 μM
6 μM
M+0
M+1
E
NHBE
Relative mRNA abundance
IFNB1
ISG15
F
NHBE
Relative RNA abundance of SARS-CoV-2
Nsp1
N
E
G
Viral titer (10² PFU/ml)
H
Caco-2
Cell viability (%)
Compounds
J
Caco-2
Relative RNA abundance of SARS-CoV-2
I
IFN-β fold induction
Mock
SeV only
(μM)
Compounds

**Figure S6. CTPS1 inhibitors suppress SARS-CoV-2 replication.** Related to **Figure 6**

(A) Effect of **Compound 1** on IRF3 deamidation was analyzed by two-dimensional gel electrophoresis and immunoblotting in A549 and LoVo cells expressing SARS-CoV-2 Nsp8.

(B) The mRNA abundance of *IFNB1* was determined by real-time PCR analysis using total RNA extracted from 293T cells pre-treated with DON, **Compound 1** or **Compound 2** for 2 h, followed with SeV infection for 9 h.

(C) Effect of **Compound 1** on NF-κB promoter activity was determined by luciferase reporter assay using 293T cells transfected with NF-κB reporter plasmid cocktail, followed with **Compound 1** treatment at indicated concentration.

(D) Effect of **Compound 1** on intracellular CTP and CDP traced with [$^{15}$N]glutamine was determined by mass spectrometry using SARS-CoV-2 ORF8-expressing LoVo cells treated with increasing amounts of **Compound 1**. M+2 is below the detection limit.

(E - G) NHBE cells with treated with **Compound 1** at indicated concentration. The mRNA abundance of *IFNB1* and *ISG15* was determined by real-time PCR analysis at 24 h after SARS-CoV-2 (MOI = 0.5) infection (E). Effect of **Compound 1** on expression of viral genes was analyzed by real-time PCR of total RNA extracted at 48 h after SARS-CoV-2 infection (MOI =0.1) (F). Medium of NHBE cells infected with SARS-CoV-2 was used for plaque assay to determine viral replication (G).

(H) Caco-2 cells were treated with **Compound 1** and its derivatives at indicated concentrations for each 24 h. Cell viability was determined by trypan blue staining at 72 h.

(I) Effects of **Compound 1** and its derivatives on IFN-β promoter activity were determined by luciferase reporter assay in 293T cells treated with the compounds and infected with SeV.

(J) RNA abundance of viral genes (*Nsp1*, *N* and *E*) in Caco-2 cells treated with **Compounds 1**, **5**, **6**, **8**, **9** and **10** was determined by real-time PCR analysis at 72 h after SARS-CoV-2 (MOI = 0.1) infection.

Error bars indicate SD of technical triplicates. Statistical significance was calculated using unpaired, two-tailed Student's *t*-test. NS, no significance; **P* < 0.05; ***P* < 0.01; ****P* < 0.001.

**Table S1, related to METHODS DETAILS**

| Real-time PCR primers for human genes | | |
|---|---|---|
| Gene Target | Forward | Reverse |
| *IFNB1* | CTTTCGAAGCCTTTGCTCTG | CAGGAGAGCAATTTGGAGGA |
| *Mx1* | GGTGGTGGTCCCCAGTAATG | ACCACGTCCACAACCTTGTCT |
| *ISG15* | GTGGACAAATGCGACGAACCCC | TCGAAGGTCAGCCAGAACAG |
| *ISG56* | TCTCAGAGGAGCCTGGCTAA | TGACATCTCAATTGCTCCAG |
| *CCL5* | CCTGCTGCTTTGCCTACATTGC | ACACACTTGGCGGTTCTTTCGG |
| *CTPS1* | AGCTTGGCAGAAGCTCTGTA | CCAACTGCATCCCTAAGCAC |
| *β-actin* | GTTGTCGACGACGAGCG | GCACAGAGCCTCGCCTT |
| ChIP-qPCR primers for mouse Ifn genes | | |
| *Ifnb1* promoter | CCAGGAGCTTGAATAAAATGAA | TGCAGTGAGAATGATCTTCCTT |
| *Ifna4* promoter | ATCCCAGACACACAAGCAGAGAG | GGCTGTGGGTTTGAGTCTTCT |
| q-PCR primers for mouse genes | | |
| *Ifnb1* | CCCTATGGAGATGACGGAGA | CCCAGTGCTGGAGAAATTGT |
| *Ifna4* | GCAGAAGTCTGGAGAGCCCTC | TGAGATGCAGTGTTCTGGTCC |
| *Isg15* | TCCATGACGGTGTCAGAACT | GACCCAGACTGGAAAGGGTA |
| *Cxcl10* | CCTGCCCACGTGTTGAGAT | TGATGGTCTTAGATTCCGGATTC |
| *Mx1* | GTGGTAGTCCCCAGCAATGT | TGCTGACCTCTGCACTTGAC |
| *Isg56* | CAAGGCAGGTTTCTGAGGAG | GACCTGGTCACCATCAGCAT |
| *Gapdh* | GTTGTCTCCTGCGACTTC | GGTGGTCCAGGGTTTCTTA |
| Real-time PCR primers for SARS-CoV-2 genes | | |
| *Nsp1* | ACACGTCCAACTCAGTTTGC | CGAGCATCCGAACGTTTGAT |
| *E* | ACTTCTTTTTCTTGCTTTCGTGGT | GCAGCAGTACGCACACAATC |
| *N* | GGGGAACTTCTCCTGCTAGAAT | GGGGAACTTCTCCTGCTAGAAT |
| sgRNA primers for Human IRF3 | | |
| IRF3 gRNA1 | CTGGTGCATATGTTCCCGGGAGG | |
| IRF3 gRNA2 | GCCGTAGGCCGTGCTTCCAAGGG | |
